# A mitochondrial electron transport chain with atypical subunit composition confers oxygen sensitivity to a mammalian chemoreceptor

**DOI:** 10.1101/2021.10.04.463079

**Authors:** Alba Timón-Gómez, Alexandra L. Scharr, Nicholas Y. Wong, Erwin Ni, Arijit Roy, Min Liu, Julisia Chau, Jack L. Lampert, Homza Hireed, Noah S. Kim, Masood Jan, Alexander R. Gupta, Ryan W. Day, James M. Gardner, Richard J. A. Wilson, Antoni Barrientos, Andy J. Chang

**Author notes:** equal contribution (Alba Timón-Gómez, Alexandra L. Scharr, Nicholas Y. Wong).

## Abstract

The carotid body (CB) is the major chemoreceptor for blood oxygen in the control of ventilation in mammals, contributing to physiological adaptation to high altitude, pregnancy, and exercise, and its hyperactivity is linked to chronic conditions such as sleep-disorder breathing, hypertension, chronic heart failure, airway constriction, and metabolic syndrome (*1–3*). Upon acute hypoxia (PO_2_=100 mmHg to <80 mmHg), K^+^ channels on CB glomus cells are inhibited, causing membrane depolarization to trigger Ca^+2^ influx and neurotransmitter release that stimulates afferent nerves (*1–3*). A longstanding model proposes that the CB senses hypoxia through atypical mitochondrial electron transport chain (ETC) metabolism that is more sensitive to decreases in oxygen than other tissues. This model is supported by observations that ETC inhibition by pharmacology and gene knockout activates CB sensory activity and that smaller decreases in oxygen concentration inhibit ETC activity in CB cells compared to other cells (*1–5*). Determining the composition of atypical ETC subunits in the CB and their specific activities is essential to delineate molecular mechanisms underlying the mitochondrial hypothesis of oxygen sensing. Here, we identify HIGD1C, a novel hypoxia inducible gene domain factor isoform, as an ETC Complex IV (CIV) protein highly and selectively expressed in glomus cells that mediates acute oxygen sensing by the CB. We demonstrate that HIGD1C negatively regulates oxygen consumption by CIV and acts with the hypoxia-induced CIV subunit COX4I2 to enhance the sensitivity of CIV to hypoxia, constituting an important component of mitochondrial oxygen sensing in the CB. Determining how HIGD1C and other atypical CIV proteins expressed in the CB work together to confer exquisite oxygen sensing to the ETC will help us better understand how tissue- and condition-specific CIV subunits contribute to physiological function and disease (*6*) and allow us to potentially target these proteins to treat chronic diseases characterized by CB dysfunction (*7*).

## HIGD1C is a novel mitochondrial protein expressed in CB glomus cells

Because the CB sensory activity correlates with changes in ETC activity (*1–3*), we sought to identify atypical mitochondrial proteins that may alter ETC responses to changes in oxygen availability. Using whole-genome expression data from RNAseq, we looked for genes encoding putative mitochondrial proteins that are overexpressed in the mouse CB compared to the adrenal medulla, a similar neuroendocrine tissue that is much less oxygen-sensitive than the CB in the adult (*8*). We found that three such genes, *Higd1c, Cox4i2, and Ndufa4l2*, were expressed at significantly higher levels in the mouse CB (Fig. 1a). By RT-qPCR, we observed similar upregulation of these genes in the human CB (Fig. 1b). In the mouse CB, *Cox4i2*, *Ndufa4l2*, and *Cox8b* were previously identified as genes upregulated by *Hif2a*, a hypoxia-inducible transcription factor critical for CB development and function (*4, 9*). Of these genes, *Cox4i2* and *Ndufa4l2* were overexpressed in the CB versus the adrenal medulla (Fig. 1a). We focused this study on *Higd1c*, the most differentially expressed of these genes and one of the top ten most upregulated genes genome-wide in the mouse CB versus adrenal medulla (*8*).

**Figure 1.**
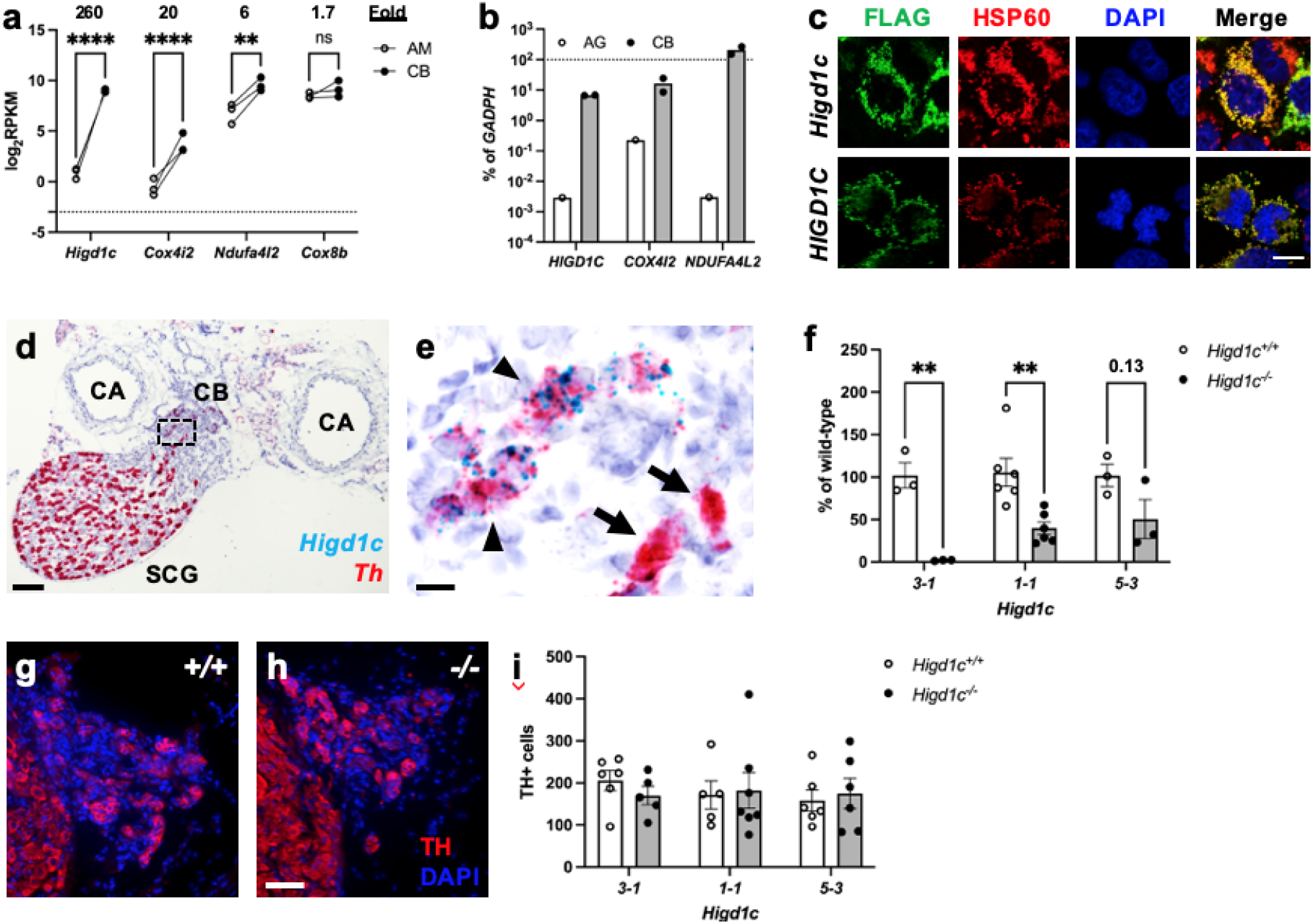
*Higd1c* expression in carotid body glomus cells is reduced in *Higd1c* CRISPR mutants. a, Expression of genes encoding four atypical mitochondrial electron transport chain (ETC) subunits in the carotid body (CB) versus adrenal medulla (AM) (*8*). RPKM, reads per kilobase per million reads mapped. n=3 cohorts of 10 animals each. Data as mean ± SEM. **p<0.01, ****p<0.0001 by two-way ANOVA with Sidak correction. b, Expression of atypical ETC proteins in human CB and adrengal gland (AG). AG, one RNA sample of adrenal glands pooled from 62 males and females, 15-61 years old. CB, two RNA samples of CBs from one male and one female adult. Dotted line, 100% of *GAPDH* expression. Data as mean. c, FLAG-tagged mouse and human HIGD1C (green) overexpressed in HEK293T cells co-localizes with the mitochondrial marker HSP60 (red) by immunostaining. DAPI, nuclear marker. Scale bar, 10 µm. d, e, BaseScope *in situ* hybridization of a wild-type C57BL/6J carotid bifurcation. e, Boxed region from d. CB, carotid body. SCG, superior cervical ganglion. CA, internal and external carotid arteries. Arrowheads, glomus cells. Arrows, SCG neurons. Scale bar, 100 µm (d), 10 µm (e). f, Expression of *Higd1c* mRNA is reduced in CBs from *Higd1c* mutants measured by RT-qPCR. n=3 (*3–1, 5–3*) and 6 (*1–1*) samples. Each sample was prepared from 4 CBs from 2 animals. Data as mean ± SEM. **p<0.01 by two-way ANOVA with Sidak correction. g, h, Immunostaining of CB glomus cells. TH, tyrosine hydroxylase. DAPI, nuclear marker. Scale bar, 50 µm. i, Quantitation of TH+ cells found no significant differences between CBs from *Higd1c^+/+^* and *Higd1c^-/-^* animals of each allele or between alleles by two-way ANOVA with Sidak correction. n=5-7 CBs from 3-7 animals. Data as mean ± SEM.

*Higd1c* is a novel member of the HIG1 hypoxia inducible domain gene family that also includes *Higd1a*, *Higd1b*, and *Higd2a*. *Higd1a* and *Higd2a*, the mammalian orthologs of the yeast respiratory supercomplex factors 1 and 2 (*Rcf1* and *Rcf2*), encode mitochondrial proteins that promote the biogenesis of ETC complexes and their assembly into supercomplexes (*10*). To determine the subcellular localization of HIGD1C, we overexpressed FLAG-tagged HIGD1C in HEK293T cells and observed that it co-localizes with the mitochondrial marker HSP60, suggesting that HIGD1C is targeted to mitochondria like HIGD1A and HIGD2A (Fig. 1c).

*Higd1a* and *Higd2a* are expressed in many mouse and human tissues and at much higher levels than *Higd1c* (*11, 12*). The ENCODE project reports very low expression of *Higd1c* in mouse tissues, except in the kidney (*13*). We found that *Higd1c* is expressed at high levels in the CB compared to other tissues in the mouse (Fig. S1a, Fig. S2a-d). Within the CB, glomus cells sense hypoxia to stimulate afferent nerves to increase ventilation (*14*). mRNA for *Higd1c* was localized in the same cells as mRNA for *Th*, a marker of glomus cells (Fig. 1d, e, Fig. S3a-l), agreeing with single-cell RNAseq findings (*15*) (Fig. S2e). As a positive control, we confirmed the previously reported expression of *Higd1c* in kidney proximal tubules (*16*) (Fig. S4a-m). These results indicate that *Higd1c* is expressed in a key cell population for oxygen sensing in the CB.

To determine if HIGD1C plays a role in CB sensory signaling, we generated mutants in *Higd1c* by CRISPR/Cas9 in C57BL/6J mice. We isolated mutants that carried large deletions that span upstream sequences through the first coding exon and small indels in the first coding exon (Fig. S1a-c). We characterized one allele representing each class of mutation (*3-1*, *1-1*, and *5-3*) that either prevent mRNA production or alter the protein sequence and cause premature stops (Fig. S1d). For all three alleles, heterozygous mutants were fertile, and homozygous mutants were viable and not underrepresented in the progeny (Table S1).

For the large deletion allele *3-1*, RT-qPCR using primers that target *Higd1c* mRNA sequences deleted in this strain amplified ∼100-fold less product from mRNA from *Higd1c^-/-^* CBs and kidneys as expected (Fig. 1f, Fig. S2a-d, Fig. S4q), confirming the specificity of this primer set. Similarly, the loss of *Higd1c in situ* hybridization signal in CBs and kidneys from *Higd1c 3-1^-/-^* mutants validated the specificity of probes for *Higd1c* exons 1 and 2 (Fig. S3m-o, Fig. S4n-p). In *1-1* and *5-3* alleles that carry mutations in exon 3, *Higd1c* mRNA levels were reduced in CBs and kidneys from *Higd1c^-/-^* mutants compared to *Higd1c^+/+^* animals (Fig. 1f, Fig. S4q). From these results, we infer that *Higd1c 3-1*, *1-1*, and *5-3* alleles all reduce HIGD1C activity.

## HIGD1C mediates CB sensory and metabolic responses to hypoxia

To assess if HIGD1C plays a role in CB oxygen sensing at the whole animal level, we performed whole-body plethysmography on awake, unanesthetized mice. A decrease in arterial blood oxygen stimulates the CB to signal to the brainstem to increase ventilation within seconds (*1–3*). *Higd1c^-/-^* mutants of all three alleles had normal ventilation in normoxia but were defective in the hypoxic ventilatory response (Fig. 2a-d, Fig. S5a-h). By contrast, *Higd1c^-/-^* mutants had robust ventilatory responses to hypercapnia comparable to *Higd1c^+/+^* animals (Fig. 2e-h, Fig. S5i-p). Because the CB is the major chemoreceptor for oxygen but only a minor contributor to CO_2_/H^+^ sensing in the control of ventilation (*14*), these results suggest that *Higd1c* specifically regulates ventilatory responses to hypoxia. The reduction in hypoxic ventilatory response in *Higd1c* mutants is most likely due to defects in the CB because *Higd1c* was expressed at low levels in the petrosal ganglion and brainstem downstream of the CB in the neuronal circuit (Fig. S2a, b).

**Figure 2.**
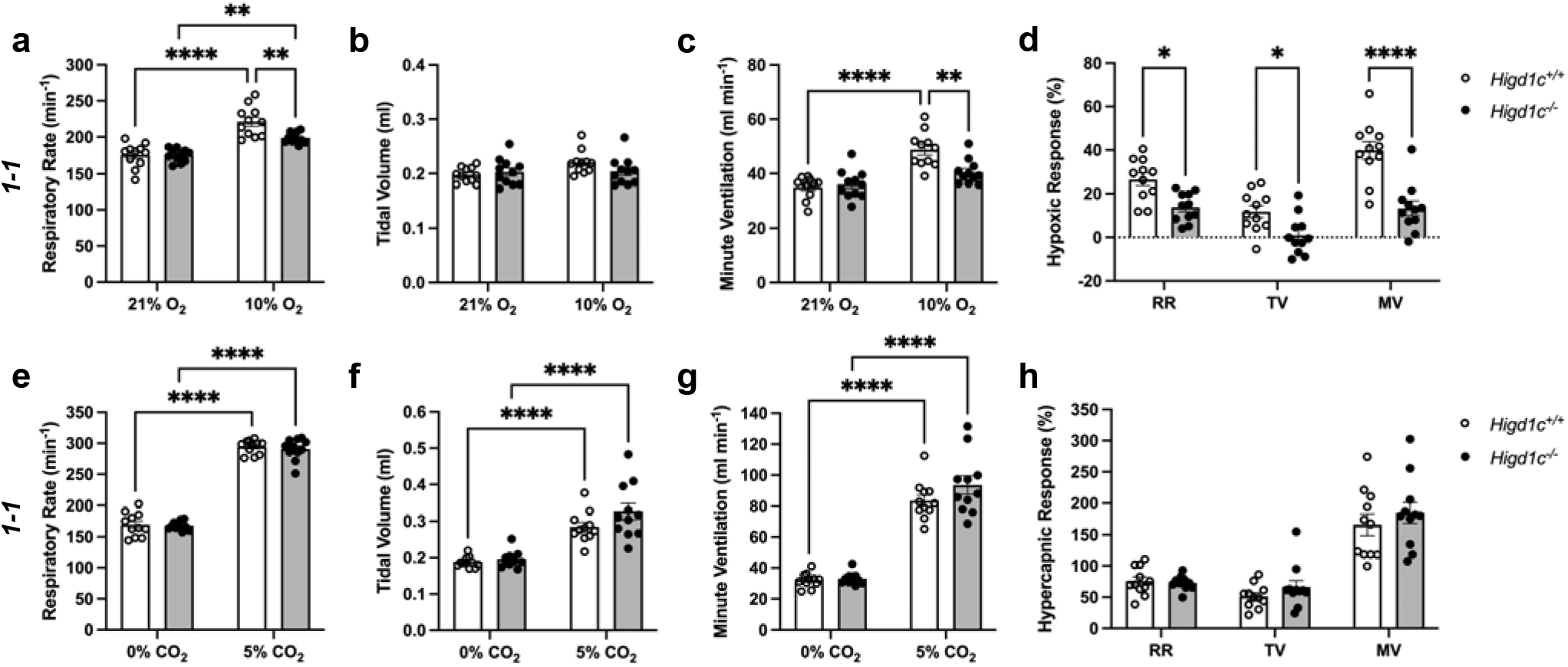
Ventilatory responses of *Higd1c* mutants to hypoxia and hypercapnia. a-h, Respiratory rate (RR), tidal volume (TV), and minute ventilation (MV) (minute ventilation = respiratory rate × tidal volume) by whole body plethysmography of unrestrained, unanesthetized *Higd1c 1-1^+/+^* and *Higd1c 1-1^-/-^* animals exposed to hypoxia (a-d) or hypercapnia (e-h). d, Hypoxic response as percentage change in hypoxia (10% O_2_) versus control (21% O_2_). h, Hypercapnic response as percentage change in hypercapnia (5% CO_2_) versus control (0% CO_2_). Ventilatory parameters of *Higd1c^+/+^* and *Higd1c^-/-^* animals in normal air conditions (21% O_2_ or 0% CO_2_) are not significantly different (p>0.05). n=11 (*+/+*), 11 (*-/-*) animals. Data as mean ± SEM. *p<0.05, **p<0.01, ***p<0.001, ****p<0.0001 by two-way ANOVA with Tukey’s test (a-c, e-g) or with Sidak correction (d, h).

Loss of function mutations in the ETC CII succinate dehydrogenase D subunit (*Sdhd*) and the transcriptional factor *Hif2a* that regulates the expression of multiple ETC subunits cause loss of CB glomus cells during development (*4, 9, 17*). The number of TH-positive glomus cells in CBs from *Higd1c^+/+^* and *Higd1c^-/-^* animals was not significantly different for all three alleles (Fig. 1g-i), and there were no gross morphological abnormalities in mutant CBs. Thus, it is unlikely that the hypoxic ventilatory response defect observed in *Higd1c^-/-^* mutants is due to the loss of glomus cells.

We next examined the integrated sensory output from the CB at the level of the carotid sinus nerve (CSN), a branch of the glossopharyngeal nerve that connects the CB to the brainstem. When the CB is activated, glomus cells release neurotransmitters to stimulate the CSN to transduce signals to the brainstem (*14*). Baseline CSN activity was similar between *Higd1c 1-1^+/+^* and *Higd1c 1-1^-/-^* tissue (Fig. 3a). As oxygen levels were decreased, CSN activity increased in a dose-dependent manner in *Higd1c 1-1^+/+^* tissue (Fig. 3b, Fig. S6a). However, this response to hypoxia was significantly attenuated in CSNs from *Higd1c 1-1^-/-^* mutants, while the response to high CO_2_/H^+^ was unaffected (Fig. 3b, Fig. S6b). Thus, we conclude that HIGD1C specifically mediates oxygen sensing at the whole organ level.

**Figure 3.**
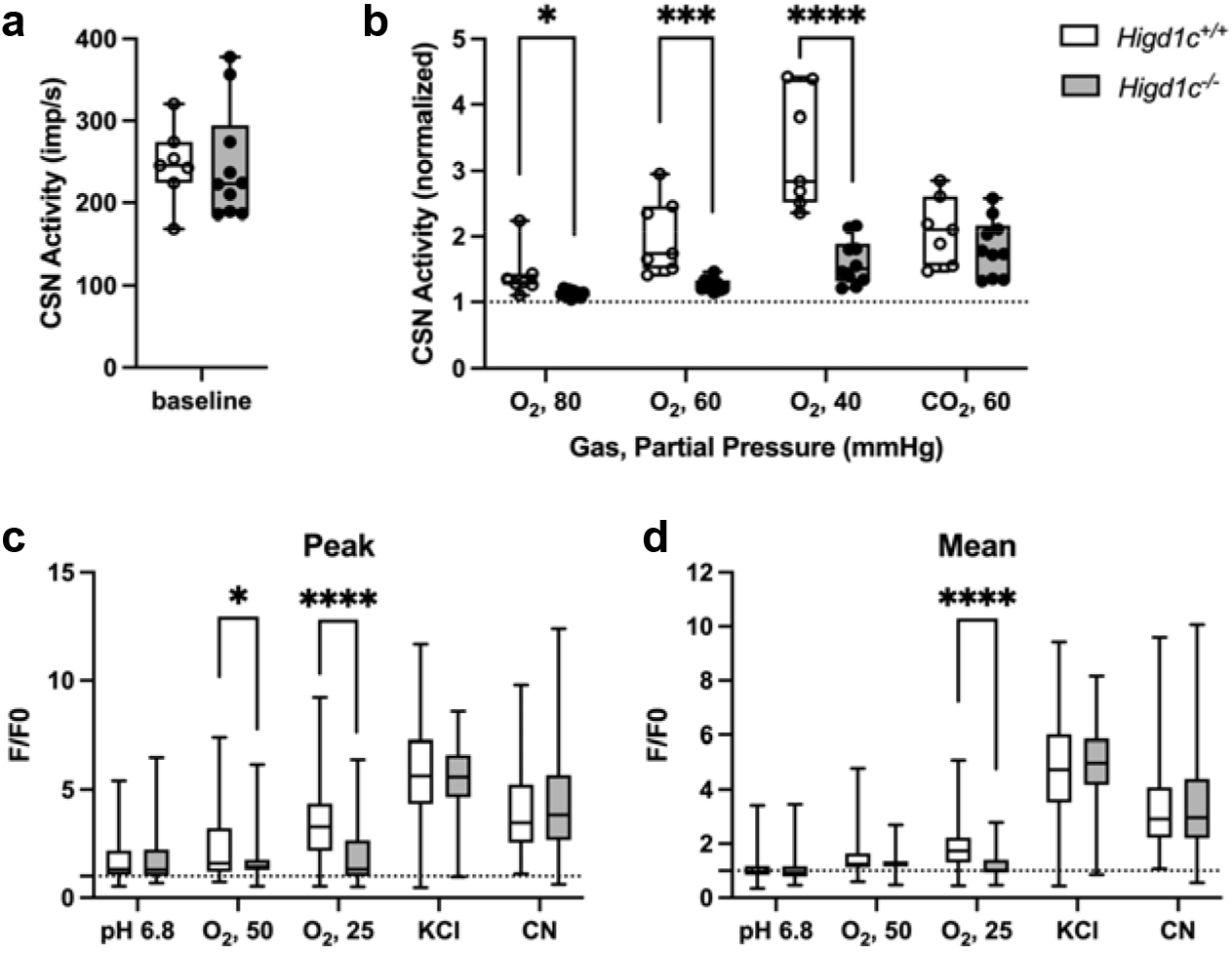
*Higd1c* mutants have defects in carotid body sensory responses to hypoxia. a, b, Quantification of carotid sinus nerve activity in *Higd1c 1-1^+/+^* and *Higd1c 1-1^-/-^* tissue preparations at baseline (a) and in hypoxia (PO_2_=80, 60, and 40 mmHg) or hypercapnia (PCO_2_=60 mmHg) (b). n=7 (*+/+*), 10 (*-/-*) preparations from 5 and 6 animals, respectively. Data as box plots showing median and interquartile interval. **p<0.01, ***p<0.001, ****p<0.0001 by Mann-Whitney U test with Holm-Sidak correction. d, e, Peak (d) and mean (e) GCaMP calcium responses (F/F0) of glomus cells from *Higd1c 1-1^+/+^* and *Higd1c 1-1^-/-^* animals to low pH (6.8), hypoxia (PO_2_=50 and 25 mmHg), high KCl (40 mM), and cyanide (CN, 1 mM). Data as box plots showing median, interquartile interval, and max/min. n= 296/4/3 (+/+), 201/4/3 (-/-) for pH 6.8, 312/5/4 (+/+), 214/5/4 (-/-) for all other stimuli. n as cells/CBs/animals. *p<0.05, ****p<0.0001 by Mann-Whitney U test with Holm-Sidak correction.

Because *Higd1c* is expressed in glomus cells (Fig. 1c, d), we evaluated if *Higd1c* mutants are also defective in the sensory responses of these oxygen-sensitive cells. Glomus cells exhibit acute calcium transients in response to stimuli, which can be visualized by the genetically encoded calcium indicator GCaMP3 (*8*). We found that glomus cells from *Higd1c 1-1^-/-^* mutants mounted a weaker calcium response to hypoxia than those from *Higd1c 1-1^+/+^* animals, with fewer glomus cells responding strongly to both levels of hypoxia (Fig. 3c, d, Fig. S6c-h). In contrast to hypoxia, glomus cells from *Higd1c 1-1^-/-^* mutants were not significantly different in their calcium response to low pH or high KCl compared to glomus cells from *Higd1c 1-1^-/-^* animals (Fig. 3d-e, Fig. S6c, d). Low pH and high KCl modulate the activity of K^+^ channels on the plasma membrane of glomus cells thought to act downstream of mitochondria in CB oxygen sensing (*18, 19*). Calcium responses to cyanide, a potent ETC CIV inhibitor, were similar between *Higd1c 1-1^+/+^* and *Higd1c 1-1^-/-^* glomus cells (Fig. 3c, d, Fig. S6c, d), suggesting that strong ETC inhibition can still trigger sensory responses in *Higd1c 1-1^-/-^* glomus cells like other mutants with defects in CB oxygen sensing (*20, 21*). Together, these results show that HIGD1C contributes specifically to glomus cell responses to hypoxia involved in oxygen sensing.

To determine if HIGD1C modulates oxygen sensitivity of mitochondria in glomus cells, we used rhodamine 123 (Rh123) to image CB inner mitochondrial membrane (IMM) potential generated by ETC activity. In normoxia and hyperoxia, Rh123 readily moves to the mitochondrial matrix, where it is quenched by self-aggregation. Hypoxia inhibits ETC activity, leading to a decrease in IMM potential and an increase in Rh123 fluorescence as the dye moves out of the mitochondrial matrix and disaggregates (*22*). *Higd1c 1-1^-/-^* glomus cells had an attenuated response to hypoxia and a greater percentage of cells that responded poorly to FCCP, a potent uncoupler of oxidative phosphorylation that depolarizes the IMM (Fig. 4a, b). If we only compared glomus cells that had strong FCCP response >0.2, we found that Rh123 fluorescence was greater in *Higd1c 1-1^-/-^* glomus cells in normoxia (PO_2_=100 mmHg), suggesting that the ETC was less active in polarizing the IMM. Nevertheless, the increase in fluorescence in hypoxia (PO_2_<80 mmHg) was smaller than in *Higd1c 1-1^+/+^* glomus cells (Fig. 4c, d, Fig. S7a-c). This pattern of IMM potential from normoxia to hypoxia resembles that of acute CII inhibition on CB sensory activity (*23*). As a control for hypoxic stimulus delivery, we found that vascular cells, which are less oxygen-sensitive than glomus cells, had a left-shifted IMM potential response to hypoxia as expected (Fig. S7d-f). These results demonstrate that HIGD1C enhances ETC inhibition by hypoxia in glomus cells, a response linked to CB sensory activity (*1–3*).

**Figure 4.**
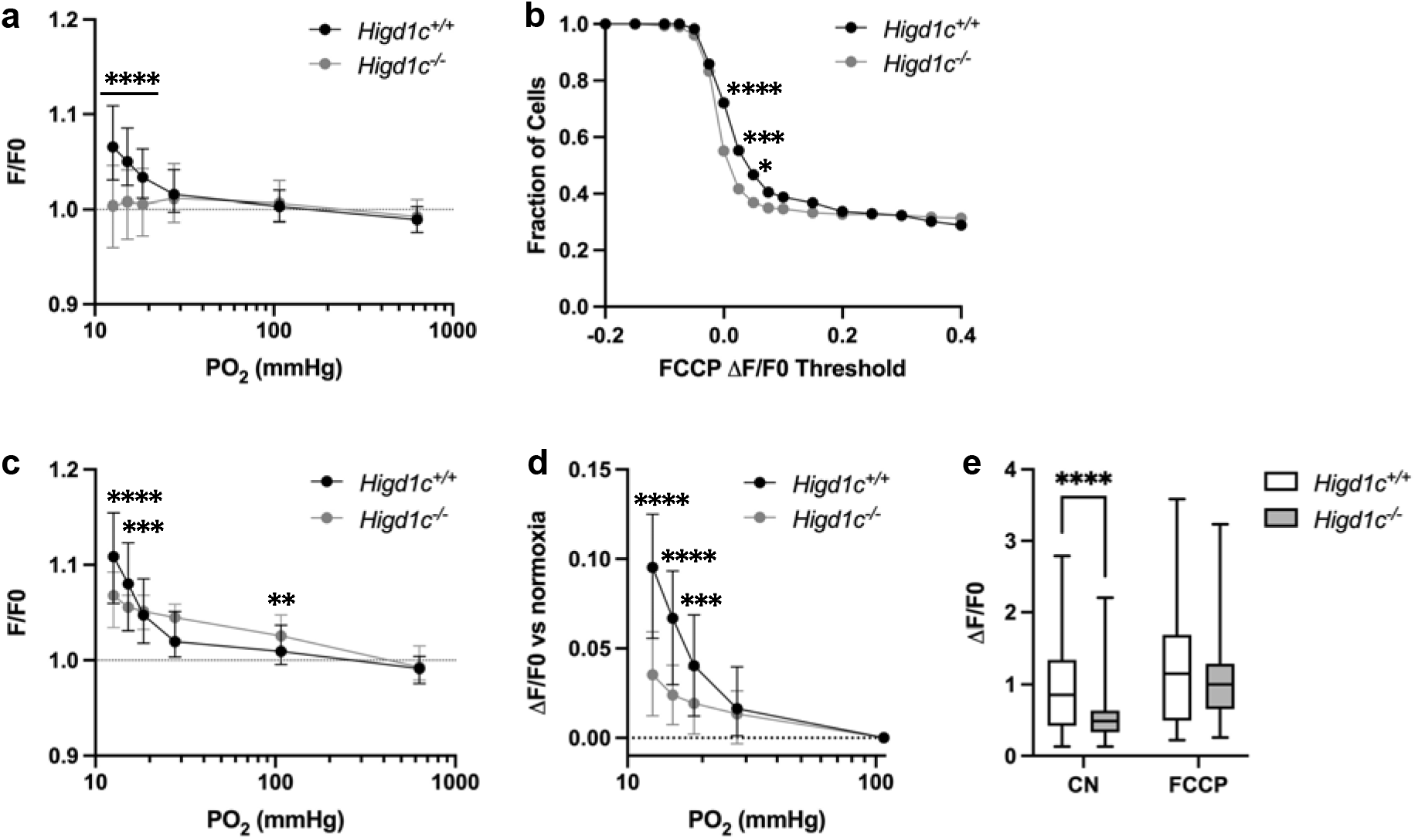
HIGD1C regulates the hypoxic response of the electron transport chain in carotid body glomus cells. a-e, Change in fluorescence of glomus cells in *Higd1c 1-1^+/+^* and *Higd1c 1-1^-/-^* CBs loaded with rhodamine 123, a dye sensitive to mitochondrial inner membrane potential, in response to hypoxia (PO_2_<80 mmHg), cyanide (1 mM), and FCCP (8 μM). Data from all glomus cells (a, b) or only glomus cells with responses to FFCP at ΔF/F>0.2 (c-e). a, c, d, Dashed line, fluorescence at the start of stimulus. a, b, n=291/3/3 (+/+), 312/3/3 (-/-) cells/CBs/animals. c-e, n=98/3/3 (+/+), 102/3/3 (-/-) cells/CBs/animals. a, c, d, Data as median and interquartile interval. e, Data as box plots showing median and interquartile interval. a, c-e, **p<0.01, ***p<0.001, ****p<0.0001 by Mann-Whitney U test with Holm-Sidak correction. b, Fraction of glomus cells that responded to FCCP at different ΔF/F. *p<0.01, ***p<0.001, ****p<0.0001 by Z test of proportions.

## HIGD1C associates with and regulates ETC Complex IV activity

To assess the role of HIGD1C in ETC function, we overexpressed FLAG-tagged human or mouse HIGD1C in HEK293T cells to perform biochemical and metabolic studies of mitochondria. In wild-type HEK293T cells, *HIGD1C* mRNA was expressed at very low levels (3×10^−6^ the level of *GAPDH* by RT-qPCR). Overexpressed FLAG-tagged HIGD1C associated with ETC CIV and cytochrome *c* (Fig. S8a-c). Notably, HIGD1C overexpression severely reduced the abundance of ETC supercomplexes, and supercomplexes that did assemble contained only traces of *in-gel* CIV activity (Fig. S8d, e). These defects in supercomplex formation correlated with a decrease in CIV enzymatic activity (Fig. S8f, g) and oxygen consumption rate (Fig. S8h, i).

Because HIGD1C is most similar to HIGD1A and HIGD2A, we overexpressed HIGD1C in *HIGD1A*-KO and *HIGD2A*-KO HEK293T cell lines to determine if it could rescue ETC defects of these KO cell lines (*24*). As in wild-type cells, HIGD1C associated with CIV and cytochrome *c* in *HIGD1A-KO* and *HIGD2A-*KO mutant cells (Fig. 5a, b, Fig. S9a-c, Fig. S10a-d). Unlike overexpression of HIGD1A, HIGD1C could not rescue defects in the assembly of supercomplexes in either *HIGD1A*-KO or *HIGD2A-*KO cells (Fig. 5c, Fig. S9d, Fig. S10e). Strikingly, however, HIGD1C overexpression restored CIV activity in *HIGD1A*-KO, but not *HIGD2A-*KO cells (Fig. 5d, Fig. S9e, f, Fig. S10f). This could be due to non-overlapping activities of HIGD1C and HIGD2A and/or more severe defects in CIV assembly in *HIGD2A*-KO cells (Fig. S9d). Instead of acting as a CIV assembly factor as HIGD2A, HIGD1C could play a regulatory role in modulating CIV activity like HIGD1A (*24*). In agreement, at the overall ETC level, cellular respiration was also restored by HIGD1C overexpression in *HIGD1A*-KO cells (Fig. S10g, h). Overexpression of mouse HIGD1C induced a weaker rescue of CIV activity than human HIGD1C, likely due to disruption of species-specific associations with ETC subunits (Fig. 5, Fig. S10f). Nonetheless, mouse HIGD1C fully rescued mitochondrial oxygen consumption due to the spare respiratory capacity of the ETC (Fig. S10g, h). These rescue experiments show that HIGD1C is not involved in ETC complex or supercomplex biogenesis, but, similarly to HIGD1A, it can interact with CIV to regulate its activity.

**Figure 5.**
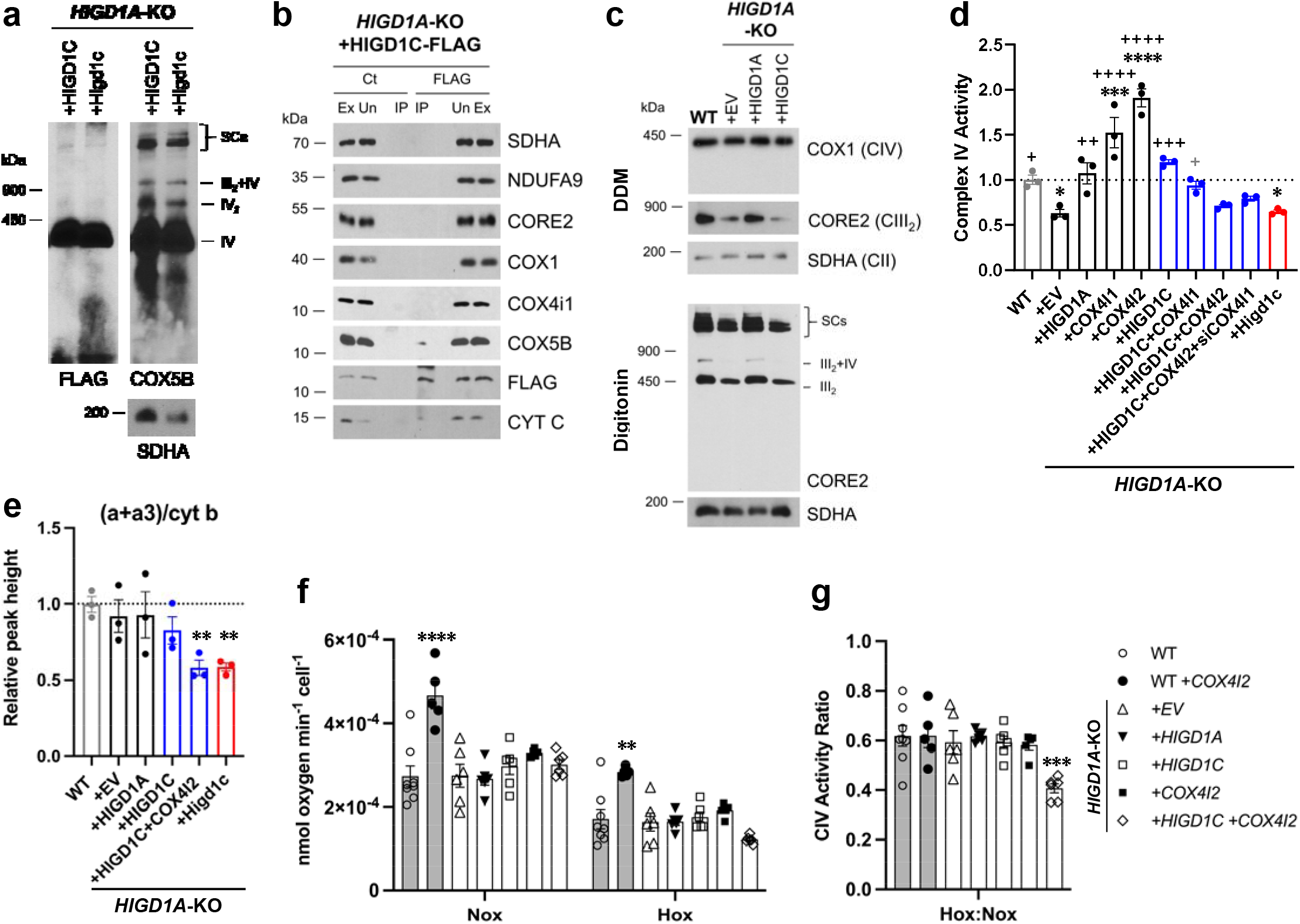
HIGD1C is a mitochondrial protein that associates with the electron transport chain Complex IV and regulates cellular respiration. a-g, *HIGD1A*-KO HEK293T cells overexpressing FLAG-tagged human or mouse HIGD1C and/or COX4 isoforms. EV, empty vector. HIGD1C, human HIGD1C. Higd1c, mouse HIGD1C. a, BN-PAGE and immunoblots using antibodies for FLAG and the Complex IV subunit COX5B. SDHA is used as a loading control. b, Co-immunoprecipitation using a FLAG antibody followed by SDS-PAGE and immunoblot using antibodies for FLAG and subunits of Complex I (NDUFA9), Complex II (SDHA), Complex III (CORE2, UQCRB), Complex IV (COX1, COX4I1, COX5B), and cytochrome *c*. c, ETC complexes and supercomplexes extracted with DDM and digitonin, respectively, detected by BN-PAGE and immunoblotting. d, Complex IV enzymatic activity assay. n=3. Data as mean ± SEM. *p<0.05, **p<0.01, ***p<0.001, ****p<0.0001 by one-way ANOVA with Dunnett test. *p-value versus WT, +p-value versus *HIGD1A*-KO+EV. Grey color indicates p=0.06. e, Relative peak height of heme *a+a3* normalized by cytochrome b peak. n=3. Data as mean ± SEM. **p<0.01 by one-way ANOVA with Dunnett test. f, Ascorbate/TMPD-dependent oxygen consumption in normoxia (Nox, PO_2_=150mmHg) and hypoxia (Hox, PO_2_=25mmHg) by high-resolution respirometry. Values normalized by cell number. n=6. g, Ratio of hypoxic/normoxic oxygen consumption calculated in f. f-g, Data as mean ± SEM. **p<0.01, ***p<0.001, ****p<0.0001 by two-way ANOVA with Sidak correction.

HIGD1C could modulate CIV activity by (1) mediating the formation of an electron-transfer bridge between ETC CIII and IV and/or (2) by producing a structural change around the active center of the enzyme. The former possibility is unlikely because overexpression of HIGD1C in *HIGD1A-*KO cells did not increase the levels of cytochrome *c* present in ETC supercomplexes (Fig. S10e) compared to control cell lines. The CIV active center is located in subunit 1 (COX1) and formed by a binuclear heme-copper center (heme *a*_3_-Cu_B_). During catalysis, reduced cytochrome *c* delivers electrons to CIV subunit 2 copper site (COX2-Cu_A_), which are then transferred to the heme *a* center in COX1, and subsequently to the active site where oxygen binds and is reduced to water (*25*). To analyze the protein environment surrounding the CIV active center, we measured UV/Vis absorption spectra of total cytochromes extracted from purified mitochondria. The absence of HIGD1A produced a blue shift in the peak of heme *a+a3* absorbance from 603 nm to 599 nm (Fig. 5e, Fig. S11a-d) that is associated with changes in the environment around the CIV heme *a* centers (*26*). Notably, previous studies showed a similar alteration of the CIV heme *a* environment in the presence of excess recombinant HIGD1A (*27*). This spectral shift observed in the KO cell line was completely restored by the expression of HIGD1A. While the wavelength at the peak appeared to be restored by overexpression of human HIGD1C (Fig. S11a), expression of human or mouse HIGD1C generated a broader peak, probably due to the existence of a mixed population of the enzyme (Fig. S11e), only partially restoring the spectral shift. A blue shift in the wavelength at the peak was apparent in *HIGD1A*-KO cells overexpressing human HIGD1C and COX4I2 together or mouse HIGD1C alone, suggesting that COX4I2 can further modify the effect of HIGD1C overexpression. These results suggest that interaction of HIGD1C with CIV can alter the active site and activity of CIV and are consistent with observations that acute hypoxia shifts the absorbance spectra of cytochromes in the CB (*1–3*).

## HIGD1C and COX4I2 enhance the sensitivity of ETC Complex IV to hypoxia

Previous studies describe the exquisite sensitivity of mitochondria of CB glomus cells to hypoxia (*1–3*). To determine if HIGD1C can alter the sensitivity of ETC to hypoxia, we measured cellular respiration of *HIGD1A*-KO cells overexpressing HIGD1C. In this cell line, the pattern of HIGD protein expression mimicked that found in mouse and human CBs, where *Higd1c* and *Higd2a* are expressed at high levels and *Higd1a* at levels undetectable by single-cell RNAseq (*15*) (Fig. S2e, f). Using an artificial electron donor system to isolate cytochrome *c*-CIV activity, we measured oxygen consumption of *HIGD1A*-KO cell lines overexpressing atypical mitochondrial subunits found in glomus cells in normoxia and hypoxia by high-resolution respirometry. Co-overexpression of HIGD1C and COX4I2 in *HIGD1A*-KO cells, which better models the ETC composition in CB glomus cells (Fig. S2e, f), decreased CIV-dependent respiration in hypoxia more than wild-type and other cell lines, including the one overexpressing HIGD1C alone, revealing an enhanced response to hypoxia (Fig. 5f, g). Overexpression of COX4I2 in HEK293T cells increased CIV-dependent respiration in both normoxic and hypoxic conditions, probably due to an increase in CIV abundance (Fig. S10i), but the ratio between conditions remained unaltered compared to the other cell lines. These results demonstrate that co-overexpression of HIGD1C and COX4I2, two atypical mitochondrial ETC proteins expressed in CB glomus cells, can confer oxygen sensitivity to HEK293T cells that are not ordinarily responsive to mild levels of hypoxia. Therefore, the COX4I2-containing ETC CIV, and its regulation by HIGD1C, emerge as the necessary and sufficient factors driving oxygen sensitivity in CB glomus cells.

## Discussion

A role for the mitochondrial ETC in CB oxygen sensing dates back to Corneille Heymans’s experiments in the 1930’s showing CB stimulation by the CIV inhibitor CN. Subsequent measurements of NADH and IMM potential in CB glomus cells at different oxygen levels demonstrate the exquisite sensitivity of their ETC to hypoxia (*1–3*). Previous studies found that mouse knockouts in specific CI and CIV subunits exhibit defects in CB sensory and metabolic responses to hypoxia (*4, 5*), phenocopying the effect of drugs that inhibit these ETC complexes. However, it is not clear whether the phenotype observed in these mutants is due to the unique oxygen sensitivity of these subunits or their essential roles in CI and CIV function.

Using whole-genome transcriptomics, we identified HIGD1C as a novel mitochondrial CIV protein essential for oxygen sensitivity of the CB. We found that HIGD1C interacts with CIV and alters the conformation of its enzymatic active center. In the absence of HIGD1A, HIGD1C and COX4I2 overexpression increased oxygen sensitivity of CIV in HEK293T cells, and overexpression of COX4I2 increased the stability of HIGD1C (Fig. S10b). Since COX4I1, the ubiquitously expressed COX4 subunit, associates with HIGD1A in CIV assembly (*24*), the alternative COX4I2 subunit may assemble with HIGD1C. In opposition to its effect in *HIGD1A*-KO cells, COX4I2 overexpression in the presence of HIGD1A in WT cells decreases HIGD1C abundance, suggesting that HIGD1A and HIGD1C interact with CIV in the same domains (Fig. S8a).

While we demonstrate here that HIGD1C and COX4I2 are sufficient to confer detectable oxygen sensitivity to CIV in HEK293T cells, additional components will be required to reconstitute the oxygen sensitivity of CB glomus cells. Other proteins upregulated in CB glomus cells, such as the CIV subunits NDUFA4L2 and COX8B and the glycolytic enzyme PCX (*4, 8*), are attractive candidates for further study to elucidate their potential contribution to the CB molecular mechanism(s) of oxygen sensing.

There is growing appreciation that CIV contains subunits that are tissue-specific and/or regulated by development, physiological changes (hypoxia and low glucose), and diseases (cancer, ischemia/reperfusion injury, and sepsis) (*6, 10*). In addition to the CB, HIGD1C is expressed in kidney proximal tubules (*16*) (Fig. S4a-m). Compared to other nephron segments, the proximal tubules have the highest oxygen demand, exhibit greater ETC sensitivity to hypoxia, and are most susceptible to ischemia/reperfusion injury (*28*). We speculate that HIGD1C modulates ETC activity in multiple oxygen-sensitive cell types to match oxygen utilization to physiological function.

## Materials and Methods

### Animals

All experiments with animals were approved by the Institutional Animal Care and Use Committees at the University of California, San Francisco, and the University of Calgary. C57BL/6J (JAX) was used as the wild-type strain. Other mouse strains obtained from repositories were Th-Cre driver: B6.FVB(Cg)-*Tg(Th-cre)^FI172Gsat^*/Mmucd (MMRRC) (*29*) and ROSA-GCaMP3: B6;129S-*Gt(ROSA)26Sor^tm38(CAG-GCaMP3)Hze^*/J (JAX) (*30*). Adult animals of both sexes from multiple litters were used in all experiments.

*Higd1c* mutants were generated by injecting C57BL/6J embryos with *in vitro* transcribed sgRNA-1 and sgRNA-2 (10 ng µl^−1^ each) together with *Cas9* mRNA (50 ng µl^−1^) and transferring injected embryos to pseudo-pregnant CD-1 females. Six founders were born and bred to C57BL/6J animals to isolate individual mutations transmitted through the germline, and sequences around sgRNA targets were PCR amplified and sequenced to identify mutations (Fig. S1). *Higd1c* mutant lines were maintained by breeding *Higd1c^+/-^* animals to each other.

### RNA purification and RT-qPCR

For mouse CB and kidney tissue, animals were anaesthetized with isoflurane and decapitated, and tissues were dissected immediately. For all other tissues, animals were anesthetized and exsanguinated by perfusing PBS through the heart before decapitation and dissection. For human tissue, CB bifurcations were procured from research-consented, de-identified organ transplant donors through a collaboration with the UCSF VITAL Core (https://surgeryresearch.ucsf.edu/laboratories-research-centers/vital-core.aspx) -- UCSF IRB-designated as non-human subjects research specimens -- and were stored and transported in Belzer UW Cold Storage Solution (Bridge to Life) on ice. CBs were then dissected in UW Solution within 18 h after harvest. After dissection, tissues were transferred to RNA*later* solution (Qiagen) and stored at 4 °C. For CB, kidney, adrenal gland, and all neuronal tissues, tissue pieces were disrupted and homogenized in a guanidine-isothiocyanate lysis buffer (Buffer RLT, Qiagen) using a glass tissue grinder (Corning), followed by a 23 gauge needle and syringe, and purified by silica-membrane columns using the RNeasy Micro Kit (Qiagen). For heart, liver, lung, and spleen, tissue pieces were ground using a glass tissue grinders (Corning) in TRIzol (Ambion), and RNA was purified by acid guandinium thiocyanate-phenol-chloroform extraction followed by isopropanol precipitation. For cell culture, cells were pelleted and resuspended in Buffer RLT before RNA purification using columns. RNA quality was assessed by visualizing 28S and 18S rRNA by agarose gel electrophoresis, and RNA concentration was measured with a Nanodrop ND-1000 Spectrophotometer (Thermo). RNA was stored at −80 °C.

Two-step RT-qPCR was performed. First, purified total RNA was synthesized into cDNA:RNA hybrids with Maxima H Minus Reverse Transcriptase (Thermo) and primed using equal amounts of oligo(dT)15 primers (Promega) and random hexamers (Thermo). RNasin Plus RNase Inhibitor (Promega) was also added to the mixture. Next, qPCR was performed using PowerUp SYBR Green Master Mix (Applied Biosystems), at 10 µl reaction volume, following manufacturer instructions. Three technical replicates were performed for each reaction and plated in TempPlate 384-well PCR plates (USA Scientific). Sample plates were run using a QuantStudio 5 Real-Time PCR System (Applied Biosystems) using a 40-cycle amplification protocol.

QuantStudio software was used to calculate threshold cycle (Ct) values. Undetermined Ct values were set to Ct=40. Ct values were averaged for all technical replicates, and normalized to either *Actb* or to *GAPDH*, using the ΔCt method.

### Cell Culture

*HIGD1A-*KO and *HIGD2A*-KO cells were constructed in HEK293T using the TALEN technology as described in (*24*). HEK293T cells were grown in 25 mM glucose Dulbecco’s modified Eagle’s medium (DMEM, Life Technologies) supplemented with 10% fetal bovine serum (FBS), 2mM L-glutamine, 1mM sodium pyruvate, and 50 µg ml^−1^ uridine without antibiotics, at 37 °C under 5% CO_2_. For metabolic imaging involving HEK293T cells, 10 mm glass coverslips were placed into 1.96 cm^2^ wells, and coated with 0.2 mg ml^−1^ poly-D-lysine for at least two hours at RT. Two days prior to the experiment, 7.5 x 10^4^ cells in 500 µl of cell media were then seeded into each well. For hypoxia experiments, cell cultures were exposed to 1% O_2_ for up to 24 h, or as controls, to standard cell culture oxygen tension (18.6% O_2_). Experiments under controlled oxygen tensions were performed in a HypOxystation® H35 (HypOxygen) to minimize undesired oxygen reperfusion. Routinely, cells were analyzed for mycoplasma contamination.

### Plasmid generation and transfection

*HIGD1C*-Myc-DDK/FLAG constructs in pCMV6-Entry were cloned under the control of a CMV promoter in the pCMV6-A-Entry-Hygro plasmid, and COX4I1/COX4I2-Myc-DDK constructs in pCMV6-Entry were cloned in the pCMV6-A-Entry-BSD, using Sfg1 and Pme1 sites. 1-2 µg of vector DNA was mixed with 5 µl of Lipofectamine™ (Thermo Fisher) in OPTIMEM-I media (GIBCO) to transfect 1.5×10^6^ cells according to manufacturer’s instructions. After 48 h, media was supplemented with 200 µg ml^−1^ of hygromycin or 10 µg ml^−1^ of blasticidin and maintained for at least 21 days.

### *In situ* hybridization

Animals were anesthetized with isoflurane, decapitated, and dissected. Tissue was fixed in RNase-free 4% PFA/PBS overnight at 4 °C and equilibrated serially in 10% sucrose/PBS for >1 h, 20% sucrose/PBS for >2 h, and 30% sucrose/PBS overnight, all at 4 °C. Tissue was then embedded in O.C.T. (TissueTek) and stored at −80 °C. The tissue was sectioned at 10 μm using a Leica CM3050S cryostat and stored in −80 °C.

Following the BaseScope protocol for fixed frozen sections, slides were baked for 50 min at 60°C and post-fixed with 10% neural-buffered saline for 15 min at 60 °C. This was followed by target retrieval for 5 min at 100 °C and protease III treatment for 30 min at 40 °C. Using the BaseScope Duplex Detection Reagent kit (Advanced Cell Diagnostics, 323810), subsequent steps of hybridization and detection followed the vendor’s protocol. Probes are listed in Table S2. The probe set for *Higd1c* was custom-designed to target only the first two exons of *Higd1c*. For detection of *Th* mRNA, amplification steps 7 and 8 were reduced from 30 min and 15 min, respectively, to 15 min and 7.5 min for some samples. Images were collected on a Nikon Ti widefield inverted microscope using a DS-Ri2 color camera.

### Immunostaining

Cultured cells on coverslips were fixed with 1% or 4% PFA/PBS for 10 min at 22 °C and used immediately or stored in PBS at 4 °C. The tissue was fixed in 4% PFA/PBS for 10 min at 22 °C and equilibrated in 30% sucrose overnight at 4 °C. Tissue was embedded in O.C.T. (TissueTek) and stored at −80 °C. Sections were cut at 10 μm using a Leica CM3050S cryostat and stored in −80 °C. Fixed cells or tissue sections were incubated with primary antibodies overnight at 4 °C. Primary antibodies were mouse anti-DDK/FLAG, mouse anti-HSP60, rabbit anti-TH, and rat anti-CD31, all used at 1:500. For kidney sections, fluorescein-labeled *Lotus tetragonolobus* lectin (LTL) was added during the primary antibody treatment. Incubation with secondary antibodies (1:250) conjugated to either Alexa Fluor 488, Alexa Fluor 555 (Life Technologies) or Cy3 (Jackson ImmunoResearch) was 45 min at 22 °C. Samples were then incubated with DAPI (1 ng ml^−1^, Life Technologies) for 5 min at 22 °C and mounted in Mowiol 4–88 (Polysciences) with DABCO (25 mg ml^−1^, Sigma-Aldrich). Samples were imaged using a Leica SPE confocal microscope for cell culture and a Zeiss Axio Observer D1 widefield inverted microscope for tissue sections.

### Whole-body plethysmography

Adult mice were removed from the housing room and placed in the procedure room for a minimum of 1 h before starting the experiment to acclimate. Ventilation of unanesthetized, awake mice was measured using a commercial system for whole-body plethysmography (Scireq). Chamber pressure was detected by a pressure transducer and temperature and humidity by a sensor. These signals were integrated using IOX2 software (Scireq) to calculate the instantaneous flow rate. Baseline breathing was established during a period of at least 30 min in control gas. The baseline was followed by two hypoxic periods and one hypercapnic period, each lasting 5 min, interspersed with recovery periods of 10 min in control gas. Gas mixtures for control, hypoxia, and hypercapnia were 21% O_2_ /79% N_2_, 10% O_2_ /90% N_2_, and 5% CO_2_ /21% O_2_ /79% N_2_, respectively (Airgas). The flow rate was held constant at 1.5 L/min.

Breathing traces were collected, and ventilatory parameters were calculated by IOX2 software (Scireq). Breath inclusion criteria were set in the software to the following: (1) inspiratory time (0.07-1 s), (2) expiratory time (0.1-1 s), (3) tidal volume (0.05-0.8 ml), and (4) respiratory rate (10-320 breaths per min). Data for all accepted breaths were exported and processed using a custom R script to calculate the average respiratory rate, tidal volume, and minute ventilation for each period. Trials were rejected if many accepted breaths occurred during periods of animal sniffing, grooming, or movement. If a trial was rejected, the animal was retested on subsequent days, for up to four trials, until stable ventilation was reached in control periods.

### Carotid sinus nerve recordings

Animals were heavily anesthetized with isoflurane and then decapitated (lower cervical level). The carotid bifurcation, including the CB, carotid sinus nerve (CSN), and superior cervical ganglion, was quickly isolated *en bloc* for *in vitro* perfusion as described previously (*31*). The carotid bifurcation was then transferred to a dissection dish containing physiological saline (115 mM NaCl, 4 mM KCl, 24 mM NaHCO_3_, 2 mM CaCl_2_, 1.25 mM NaH_2_PO_4_, 1 mM MgSO_4_, 10 mM glucose, 12 mM sucrose) bubbling 95% O_2_/5% CO_2_. After 15-20 min, the isolated tissue was transferred to a recording chamber (AR; custom made) with a built-in water-fed heating circuit, and the common carotid artery was immediately cannulated for luminal perfusion with physiological saline equilibrated with 100 mmHg PO_2_ and 35 mmHg PCO_2_ (balance N_2_). The CSN was then carefully desheathed, and the carotid sinus region was bisected. The occipital and internal and external carotid arteries were ligated, and small incisions were made on the internal and external carotid arteries to allow perfusate to exit. A peristaltic pump was used to set the perfusion rate at 7 ml min^−1^, which was sufficient to maintain a constant pressure of 90-100 mmHg at the tip of the cannula (AR, custom made). The perfusate was equilibrated with computer-controlled gas mixtures using CO_2_ and O_2_ gas analyzers (CA-2A and PA1B, Sable Systems); a gas mixture of 100 mmHg PO_2_ and 35 mmHg PCO_2_ (balance N_2_) was used to start the experiments (yielding pH ∼7.4). Before reaching the cannula, the perfusate was passed through a bubble trap and heat exchanger. The temperature of the perfusate, measured continuously as it departed the preparation, was maintained at 37 ± 0.5°C. The effluent from the chamber was recirculated.

Chemosensory discharge was recorded extracellularly from the whole desheathed CSN, which was placed on a platinum electrode and lifted into a thin film of paraffin oil. A reference electrode was placed close to the bifurcation. CSN activity was monitored using a differential AC amplifier (Model 1700, A-M Systems) and a secondary amplifier (Model 440, Brownlee Precision). The neural activity was amplified, filtered (0.3 – 1kHz), displayed on an oscilloscope, rectified, integrated (200 ms time constant), and stored on a computer using an analog-to-digital board (Digidata 1322A, Axon Instruments) and data acquisition software (Axoscope 9.0). Preparations were exposed to a brief hypoxic challenge (PO_2_=60□mmHg) to determine viability; preparations that failed to show a clear increase in activity during this challenge were discarded. After this challenge, the preparations were left undisturbed for 30□min to stabilize before the experimental protocol was initiated.

The following protocol was used for all experiments: (1) the CB was perfused for 5 min with normoxia (100 mmHg PO_2_/35 mmHg PCO_2_) to determine baseline CSN activity; (2) neural responses were obtained by challenging the carotid body for 4 min with mild, moderate, and severe hypoxia (80, 60 and 40 mmHg PO_2_, respectively) interspersed with normoxia; (3) a hypercapnic (60 mmHg PCO_2_) challenge was given for 4 min (Fig. S6a, b).

Data were analyzed offline using custom software (written by RJAW). CSN activity was divided into 60 s time bins, and the activity in each bin was rectified and summed (expressed as integrated neural discharge). The neural responses for different conditions in the protocol were normalized to the baseline (normoxic) condition. Data acquisition and CSN activity analysis were performed blinded to genotype.

### Calcium imaging

*Th-Cre^Tg/+^; ROSA-GCaMP3^Tg/Tg^; Higd1c^+/+^* and *Th-Cre^Tg/+^; ROSA-GCaMP3^Tg/Tg^; Higd1c^-/-^* animals expressing GCaMP3 in glomus cells were generated, and CB was imaged as previously described (*8*). Animals were anesthetized with isoflurane and decapitated. Carotid bifurcations were dissected and cleaned in PBS to keep only the CB attached to the bifurcation. The preparation was then incubated in a physiological buffer (115 mM NaCl, 5 mM KCl, 24 mM NaHCO_3_, 2 mM CaCl_2_, 1 mM MgCl_2_, 11 mM glucose) at 26°C in a tissue culture incubator with 5% CO_2_ before transfer to the recording chamber for imaging.

At baseline, the CB was superfused by gravity at 5 ml min^−1^ with physiological buffer bubbling 95% O_2_/5% CO_2_ in the reservoir to maintain PO_2_∼700 mmHg and pH 7.4 in the imaging chamber at 22 °C. Buffer pH was lowered to 6.8 by reducing NaHCO_3_ to 7 mM with equimolar substitution of NaCl while bubbling 95% O_2_/5% CO_2_. Two levels of hypoxia at PO_2_∼25 mmHg and 50 mmHg were generated by bubbling physiological buffer in the reservoir with 90% N_2_/5% O_2_/5% CO_2_ and 95% N_2_/5% CO_2_, respectively. The preparation was sequentially stimulated with low pH and hypoxia for periods of 4.5 min each, with 3 min of recovery between stimuli. These were followed by KCl (40 mM) and CN (1 mM) for periods of 2.25 min each, with 4.5 min of recovery between stimuli (Fig. S6c, d).

Imaging was performed on a Zeiss LSM 7 MP two-photon microscope with a Coherent Ultra II Chameleon laser and a sensitive gallium arsenide phosphide (GaAsP) detector. Preparations were excited at 960 nm, and emission was collected at 500-550 nm. Using a 20X water immersion objective, we acquired Z-stacks at 2 µm intervals at a resolution of 1024 by 1024 pixels and up to 60-85 µm of tissue depth.

Regions of interest corresponding to individual glomus cells were identified and analyzed in ImageJ. All regions of interest are included in the data. Fpre fluorescence was defined as the average fluorescence over the 4 frames immediately prior to the onset of the stimulus in the chamber. Mean and peak fluorescence was calculated over the duration when the stimulus was present in the imaging chamber. The ratio of Fstim/Fpre was calculated by dividing the mean and peak by Fpre just preceding the stimulus. Data acquisition and ROI analysis were carried out blinded to genotype.

### Metabolic imaging

*Higd1c^+/+^* and *Higd1c ^-/-^* animals were anesthetized with isoflurane and decapitated. Carotid bifurcations were dissected and cleaned in PBS to keep only the CB attached to the bifurcation. The preparation was then incubated in 50 µg ml^−1^ rhodamine 123 (ThermoFisher) in a physiological buffer (115 mM NaCl, 5 mM KCl, 24 mM NaHCO_3_, 2 mM CaCl_2_, 1 mM MgCl_2_, 11 mM glucose) at 26 °C in a tissue culture incubator with 5% CO_2_ for 30 min before transfer to the recording chamber for imaging.

At baseline, the CB was superfused by gravity at 5 ml min^−1^ with physiological buffer bubbling 95% O_2_/5% CO_2_ in the reservoir to maintain PO_2_∼700 mmHg and pH 7.4 in the imaging chamber at 22 °C. Hypoxia down to PO_2_∼10 mmHg was generated by bubbling physiological buffer in the reservoir with 95% N_2_/5% CO_2_ for 7.5 min. PO_2_ was measured using a Clark style oxygen sensor (OX-50, Unisense). After 6 min of baseline recording, the preparation was stimulated with hypoxia for a period of 7.5 min followed by 7.5 min of recovery between stimuli. This was followed by CN (1 mM) and FCCP (2 µM) for periods of 2.25 min each, with 7.5 min of recovery between stimuli.

Imaging was performed on a Zeiss LSM 7 MP two-photon microscope with a Coherent Ultra II Chameleon laser and a sensitive gallium arsenide phosphide (GaAsP) detector. Preparations were excited at 960 nm, and emission was collected at 500-550 nm. Using a 20X water immersion objective, we acquired Z-stacks at 2 µm intervals at a resolution of 1024 by 1024 pixels and up to 60-85 µm of tissue depth.

Regions of interest (ROIs) corresponding to individual glomus cells were identified and analyzed in ImageJ. All regions of interest are included in the data except as indicated in specific analyses. Due to a linear decrease in baseline fluorescence over the time course of the experiment, baseline subtraction was performed. First, the fluorescence trace of each ROI was smoothed using a 3 point centered rolling average, and the baseline was calculated using linear interpolation between the inter-stimulus intervals. This baseline was then subtracted from the original traces. Fpre fluorescence was defined as the fluorescence immediately prior to the onset of the stimulus in the chamber. Mean and peak fluorescence were calculated over the duration when the stimulus was present in the imaging chamber. The ratio of Fstim/Fpre was calculated by dividing the mean and peak by Fpre just preceding the stimulus. Data acquisition and ROI analysis were carried out blinded to genotype.

### Mitochondrial biochemistry

Mitochondrial fractions were obtained as previously described in (*24, 32, 33*) from ten 80% confluent 15-cm plates or from 1 L of liquid culture. Whole-cell extracts were obtained from pelleted cells solubilized in RIPA buffer (25 mM Tris-HCl (pH 7.6), 150 mM NaCl, 1% NP-40, 1% sodium deoxycholate, and 0.1% SDS) with 1 mM PMSF for 20 min. Extracts were cleared after 5 min centrifugation at 15,000 rpm at 4 °C.

Proteins from purified mitochondria were extracted in native conditions with either digitonin at a proportion of 1:2 of protein or with n-dodecyl-β-D-maltoside (DDM) at a concentration of 0.4%. Samples were incubated on ice for 10 min and pelleted at 10,000 x g for 30 min at 4 °C. Samples were prepared for Blue Native Electrophoresis and/or Complex I and Complex IV *in-gel* activity (IGA) assays as described (*34*). Immunoprecipitation of HIGD1C-Myc-DDK-tagged proteins was performed using 1 mg of mitochondria, extracted in 1.5 M aminocaproic acid, 50 mM Bis-Tris pH 7, 1% digitonin, 1 mM PMSF, and 8 µl of protease inhibitor cocktail (Sigma, P8340) for 10 min on ice. Samples were pelleted at 10,000 x g for 30 min at 4 °C, and the extract (Ex) was incubated for 2-3 h at 4 °C with 30 µl of FLAG-conjugated beads (anti-DYDDDDDK beads, Sigma) or empty beads (ThermoScientific), previously washed in PBS. Beads were washed 5 times in 1 mL of 1.5 M aminocaproic acid, 50 mM Bis-Tris (pH 7), 0.1% digitonin, and boiled for 5 min with 50 µl of Laemmli buffer two times to release bound material. Representative amounts of all fractions were loaded on 14% SDS-PAGE gels.

### Complex specific assay and oxygen consumption rate

Mitochondrial respiratory chain Complex IV activity was performed according to established methods (*35*). Citrate synthase activity was used as a control. Enzymatic activities were expressed relative to the total amount of extracted protein.

Oxygen consumption rate (OCR) in normoxia was measured polarographically using a Clark-type electrode from Hansatech Instruments (Norfolk, United Kingdom) at 37 °C. ∼2×10^6^ cells were trypsinized and washed with PBS, and then resuspended in 0.5 ml of permeabilized-cell respiration buffer (PRB) containing 0.3 M mannitol, 10 mM KCl, 5 mM MgCl_2_, 0.5 mM EDTA, 0.5 mM EGTA, 1 mg ml^−1^ BSA, and 10 mM KH_2_PO_4_ (pH 7.4) at 37 °C, supplemented with 10 units of hexokinase. The cell suspension was immediately placed into the polarographic chamber to measure endogenous respiration. Digitonin permeabilization (0.02 mg ml^−1^) was performed to assay substrate-driven respiration, using FADH-linked substrates (10 mM succinate plus 5 mM glycerol-3-phosphate) in the presence of 2.5 mM ADP (phosphorylation state). Oligomycin-driven ATP synthesis inhibition (0.75 µg ml^−1^) was assayed to obtain the non-phosphorylating state. Maximal oxygen consumption was reached by successive addition of the uncoupler CCCP (up to 0.4 µM). 0.8 µM KCN was used to assess the mitochondrial specificity of the oxygen consumption measured, and values were normalized by total cell number.

High-resolution respirometry was used to determine mitochondrial oxygen consumption and ascorbate/TMPD-dependent respiration in normoxia and hypoxia. Measurements were performed in intact or digitonin-permeabilized cells, respectively, in an Oxygraph-2k (Oroboros Instruments). Assays were performed according to manufacturer’s SUIT protocols, using 2-4×10^5^ cells washed with PBS and resuspended in Mir05 medium, and results were normalized by cell number.

### Extraction of total mitochondrial cytochromes

8 mg of mitochondria were extracted with 330 mM KCl, 50 mM Tris-HCl (pH 7.5), and 10% potassium deoxycholate. Samples were mixed by inversion three times and pelleted for 15 min at 40,000 x g at 4°C. The clear supernatant was transferred to a new tube, and a final concentration of 2% potassium cholate was added. The extract was divided into two equal aliquots in 1 ml quartz cuvettes, and the baseline was established. Then, the reference aliquot was oxidized with potassium ferricyanide, and the other was reduced with a few grains of sodium dithionite. Differential reduced *vs* oxidized spectrum was recorded from 450 to 650 nm.

### Data analysis and statistics

Data analysis and statistical tests were performed using Microsoft Excel, custom-written scripts in R, and GraphPad Prism software. All data are biological replicates. Group data were analyzed by the Shapiro-Wilk test to determine if data was normally distributed with the critical *W* value set at a 5% significance level. Normally distributed data are presented as mean ± standard error of the mean (SEM) and compared by two-way analysis of variance (ANOVA) followed by Tukey’s test for all pairwise comparisons, Sidak correction for multiple pairwise comparisons, or Dunnett’s test for multiple comparisons to a single group. For comparisons that included groups that did not fit the assumption of normal distribution, data are presented as box plots with median, 25^th^, and 75^th^ percentiles indicated and compared by Mann-Whitney U tests followed by Holm-Sidak correction for multiple comparisons. The Z test of proportions was used to compare the proportion of glomus cells with calcium responses at different thresholds. Chi-square test was performed to determine if the distributions of glomus cells responsive to different hypoxic stimuli were drawn from the same population. All tests were two-tailed. No statistical method was used to predetermine sample size.

## Acknowledgements

We acknowledge Alex Diaz de Arce for assistance with RNAseq analysis, Kailyss Freeman and the UCSF VITAL Core for coordinating donor tissue, ACD Bio for BaseScope *in situ* hybridization, Blair Gainous, Pauline Colombier, Brian Black, and the CVRI Transgenic Mouse Core for guidance and technical support in generating *Higd1c* mutant mice, Chris Allen for use of his two-photon microscope and cryostat, and the UCSF Nikon Imaging Center for use their widefield microscope. We sincerely thank Donor Network West, and most importantly the organ donors and their families, who give this precious gift to further scientific research.

The work was supported by a Canadian Institute of Health Research (201603PJT/366421 to R.J.A.W), a NIGMS-R35 grant (GM118141 to A.B.), and a Sandler Program for Breakthrough Biomedical Research New Frontier Award and UCSF CVRI (A.C.). A.T. was supported by a Muscular Dystrophy Association (MDA) Development Grant (MDA 862896), A.G. by an UCSF Transplant T32 FAVOR grant (P0548805), and J.G. by an UCSF Physician-Scientist Scholars Program (PSSP) grant. RJAW is an AIHS Senior Scholar.

---

**Figure S1.**
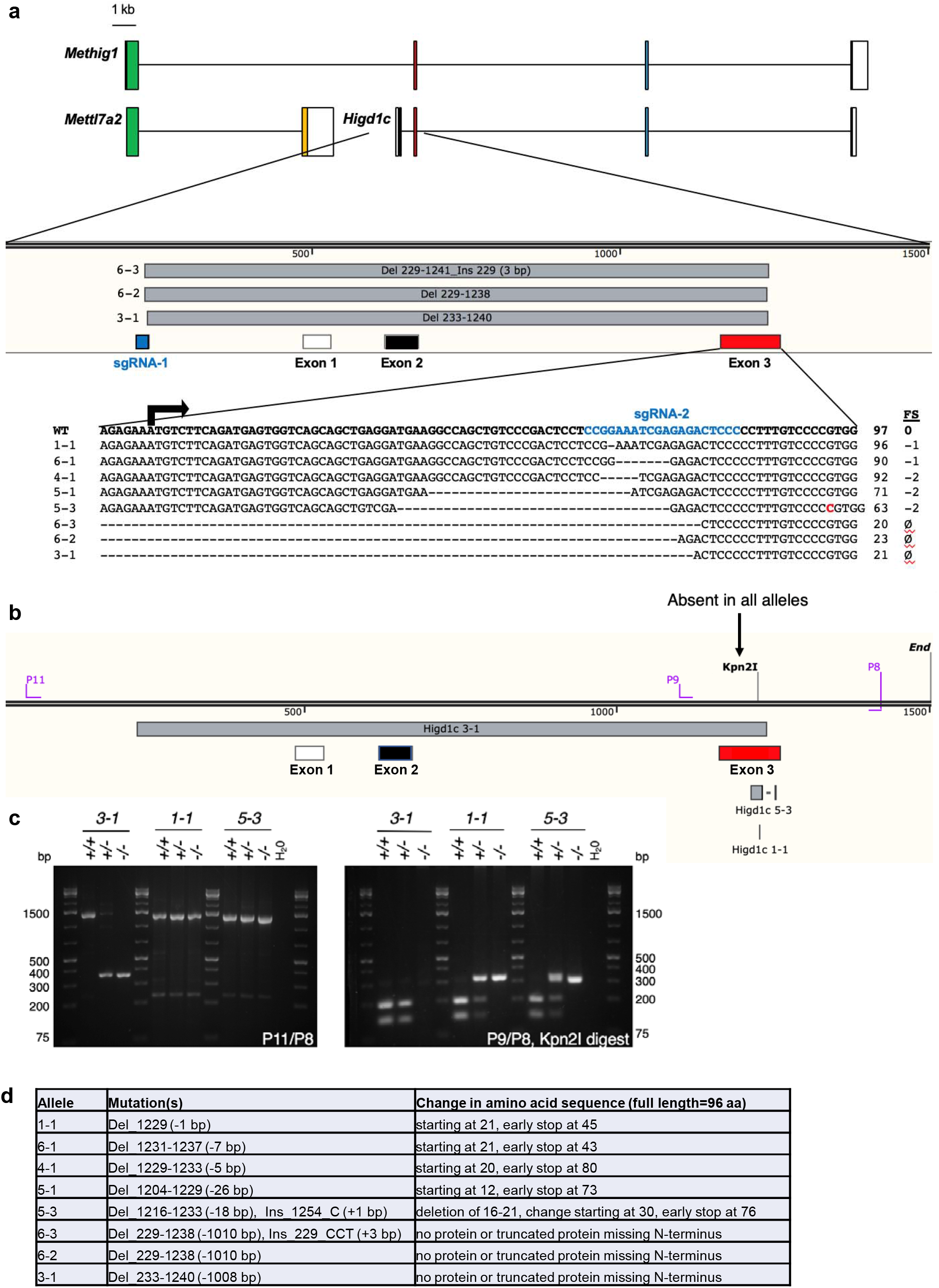
Generation and mapping of *Higd1c* CRISPR mutants. a, Schematic representation of mRNA transcripts of *Higd1c* and *Mettl7a2* and a fusion transcript spanning both genes named *Methig1*. Boxes denote exons. Arrow, *Higd1c* translational start site. Two sgRNAs targeting sequences upstream of exon 1 and within exon 3 (first coding exon) of *Higd1c* were injected together with *Cas9* mRNA into C57BL/6J embryos. Six founders were generated, from which 8 alleles were identified by sequencing around the sgRNA target sites. Mutant alleles isolated included short indels around the sgRNA target sequence in exon 3 and deletions that spanned the regions between the two guides. Arrow denotes the start codon of *Higd1c* in exon 3. FS, frameshift mutation. ø, no transcript or truncated transcript starting in exon 4. b, Genotyping primers for large deletion and small indel *Higd1c* alleles. All mapped mutations abolish a restriction site for Kpn2I in exon 3. c, PCR with P11/P8 primer pair used to genotype the large deletion 3-1 allele. PCR with P9/P8 primer pair followed by restriction digest with KpnI used to genotype small indels in exon 3. d, Expected changes to the amino acid sequence of HIGD1C in mutant alleles.

**Figure S2.**
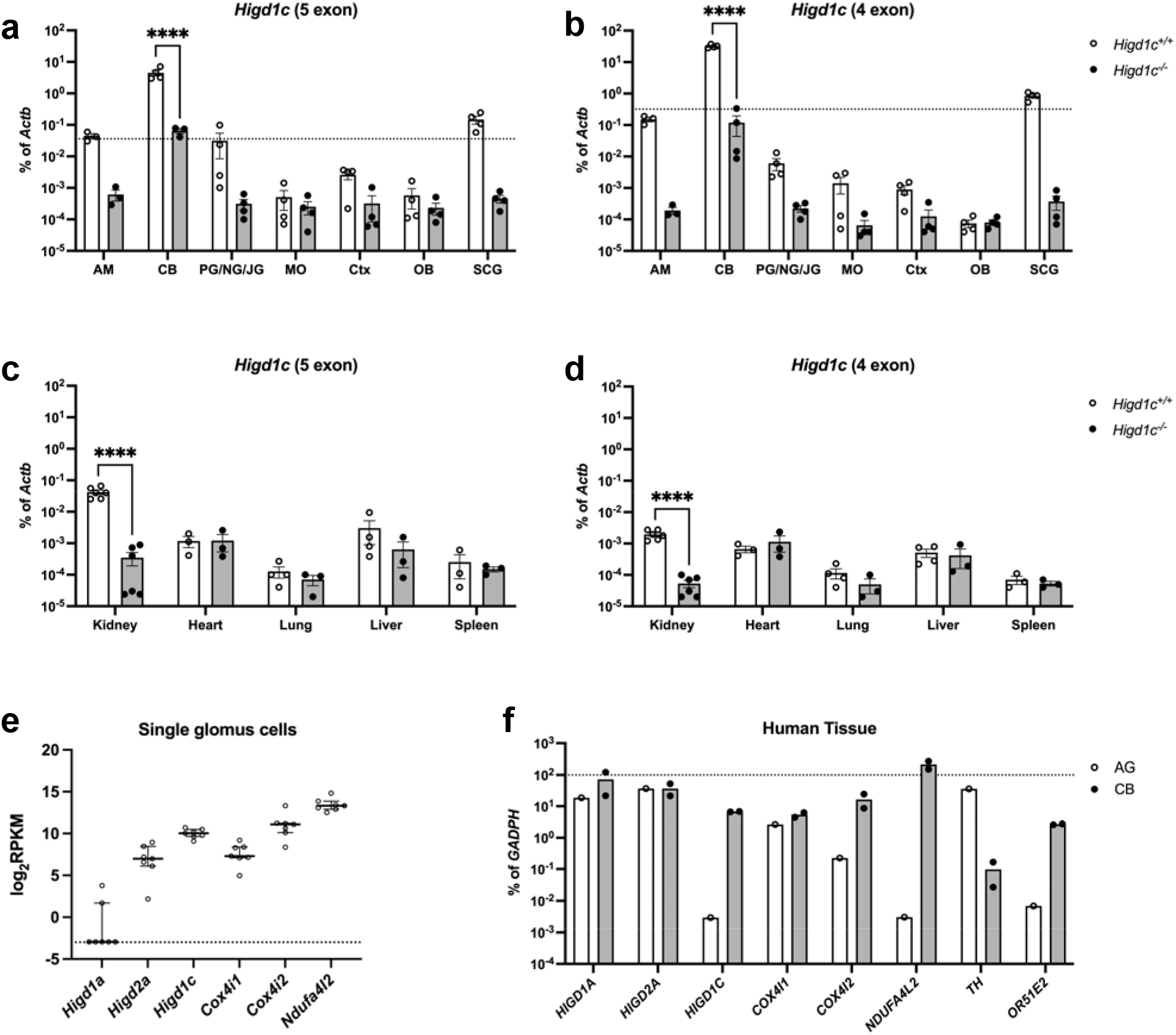
*Higd1c* is highly expressed in the CB compared to other tissues. a-d, qRT-PCR targeting *Higd1c* mRNA transcripts containing 5 exons (a, c) or 4 exons (b, d) in tissues from *Higd1c 3-1^+/+^* and *Higd1c 3-1^-/-^* animals. These mRNA isoforms differ from each other only in the first two exons that encode 5’ UTR; the 4 exon isoform does not contain Exon 2 (Fig. S1a). *Higd1c* expression normalized to *Actb* encoding *β*-actin by template and day of the experiment. Dotted line, 1% of the CB expression level. AM, adrenal medulla. CB, carotid body. PG/NG/JG, petrosal ganglion/nodose ganglion/jugular ganglion. MO, medulla oblongata. Ctx, cerebral cortex. OB, olfactory bulb. SCG, superior cervical ganglion. For CB, n=3/6 (a) and 4/8 (b) samples/animals with 4 CBs from 2 animals pooled per sample. For kidney AM, n=8/8 samples/animals (c, d). For all others, n=3/3 or 4/4 samples/animals (a-d). Data as mean and SEM. ****p<0.0001 by two-way ANOVA with Sidak correction. Because *Actb* expression can vary between tissues, we used *Higd1c 3-1^-/-^* tissues carrying a deletion that abolishes one primer target site as a negative control for *Higd1c* expression. e, Expression of select mitochondrial ETC subunits by single-cell RNAseq of 7 mouse glomus cells from a previously published dataset (*15*). One cell from this study was excluded from analysis because it had little to no expression of genes known to be expressed at high levels in mouse glomus cells, such as *Olfr78*, *Epas1*, *Cox4i2*, and *Ndufa4l2* (*4, 8*). RPKM, reads per kilobase per million reads mapped. Data presented as the median and interquartile interval. Dotted line, RPKM level corresponding to no reads. f, Expression in total RNA from human tissues of select mitochondrial ETC subunits and other genes known to be expressed in the CB. Unlike its expression in mice, the tyrosine hydroxylase gene *TH* is expressed in only a subset of glomus cells in the adult human CB (*36, 37*). The olfactory receptor gene *OR51E2* is the ortholog of mouse *Olfr78*, one of the most highly upregulated genes in the CB versus adrenal medulla in mice (*8*). AG, one RNA sample of adrenal glands pooled from 62 males and females, 15-61 years old. CB, two RNA samples of carotid bodies from two transplant donors, a 55 year-old male and a 35 year-old female. Dotted line, 100% of *GAPDH* expression. Data presented as the mean.

**Figure S3.**
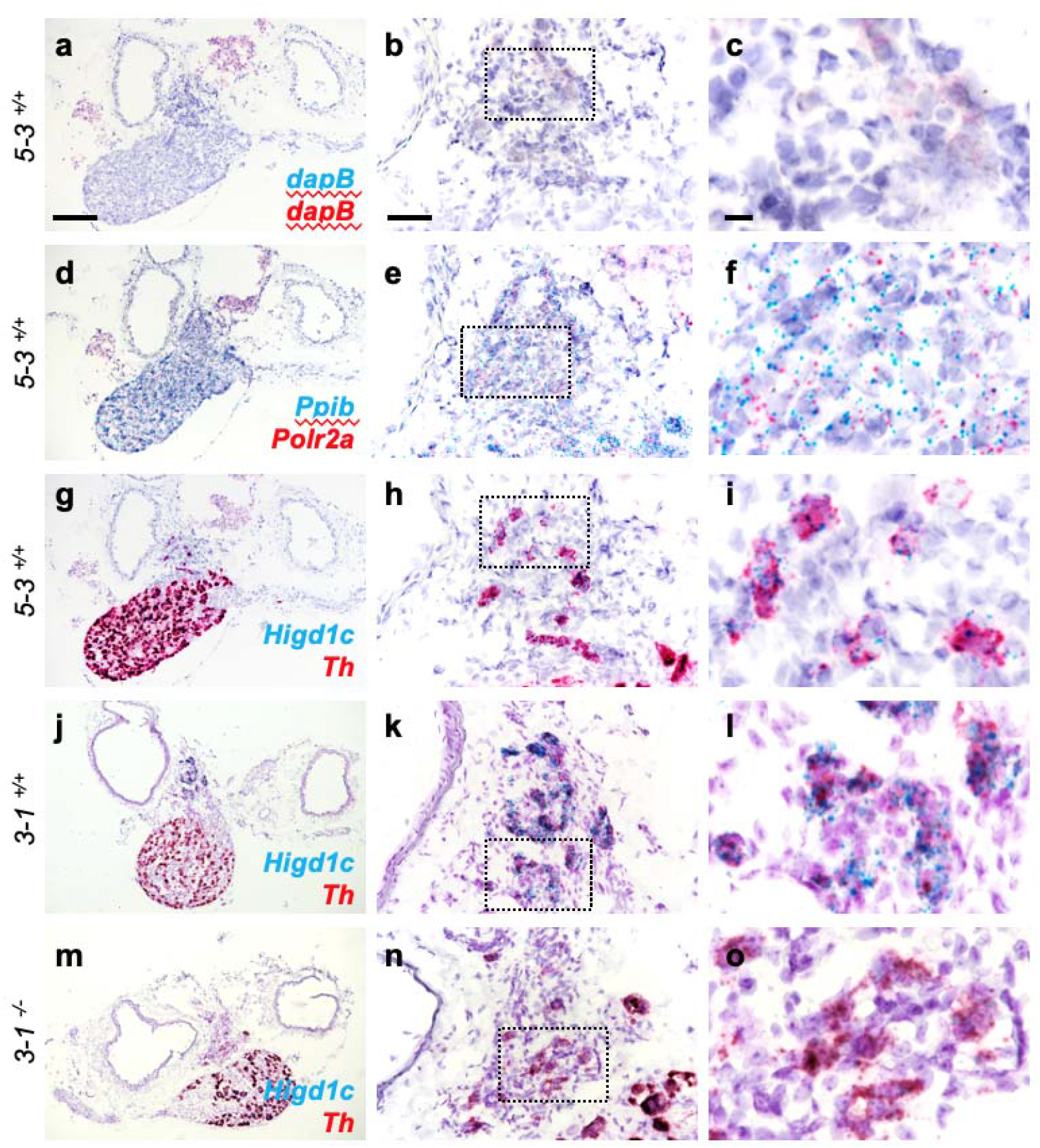
*Higd1c* is expressed in glomus cells of the carotid body. a-o, BaseScope *in situ* hybridization to detect mRNA transcripts in adjacent sections of the carotid bifurcation from a *Higd1c 5-3^+/+^* animal and from *Higd1c 3-1^+/+^* (j-l), and *Higd1c 3-1^-/-^* (m-o) animals. c, f, i, l, and o, Boxed regions from b, e, h, k, and n, respectively. a-c, *dapB* probes are negative controls that target bacterial genes. d-f, *Ppib* and *Polr2a* are positive control probes for mouse transcripts expressed ubiquitously. g-o, Probes that target transcripts for *Higd1c* and tyrosine hydroxylase (*Th*), a marker of glomus cells. The large deletion in the *Higd1c 3-1* allele removes the target sequence of BaseScope probes for *Higd1c*. Scale bars, 200 µm (a, d, g, j, and m), 50 µm (b, e, h, k, and n), and 10 µm (c, f, I, l, and o).

**Figure S4.**
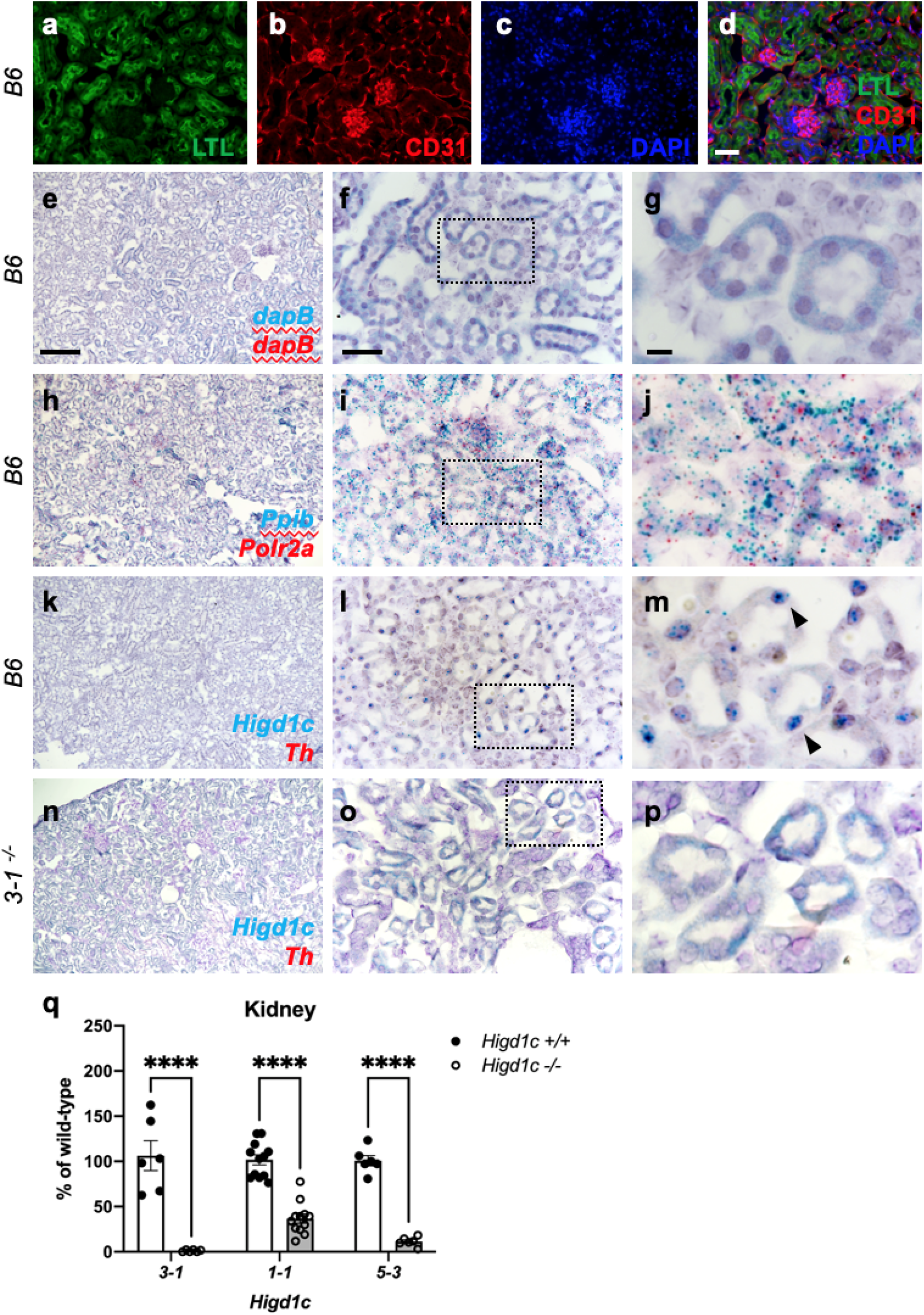
*Higd1c* expression in kidney proximal tubules is attenuated in *Higd1c* mutants. a-d, Staining of the kidney from a wild-type C57BL/6J (B6) animal with fluorescein-conjugated *Lotus tetragonolobus* lectin (LTL) that binds to the apical surface of proximal tubules and an antibody for CD31, a marker of endothelial cells. DAPI, nuclear marker. Scale bar, 50 µm. e-p, BaseScope *in situ* hybridization to detect mRNA transcripts in sections of the kidney from a wild-type C57BL/6J (B6) animal (e-m) and from a *Higd1c 3-1^-/-^* mutant (n-p). g, j, m, and p, Boxed regions from f, i, l, and o, respectively. e-g, *dapB* probes are negative controls that target bacterial genes. h-j, *Ppib* and *Polr2a* are positive control probes for mouse transcripts expressed ubiquitously. k-p, Probes that target transcripts for *Higd1c* and tyrosine hydroxylase (*Th*), a marker of glomus cells. The large deletion in the *Higd1c 3-1* allele removes the target sequence of BaseScope probes for *Higd1c*. Scale bars, 200 µm (e, h, k, and n), 50 µm (f, i, l, and o), and 10 µm (g, j, m, and p). q, Expression of *Higd1c* mRNA is reduced in kidneys from *Higd1c* mutants as measured by qRT-PCR. The target region of one primer of the primer pair is deleted in the *Higd1c 3-1* allele. n=6 (*3–1, 5–3*), 12 (*1–1*) kidneys from 6 and 12 animals, respectively. Data as mean ± SEM. ****p<0.0001 by two-way ANOVA with Sidak correction.

**Figure S5.**
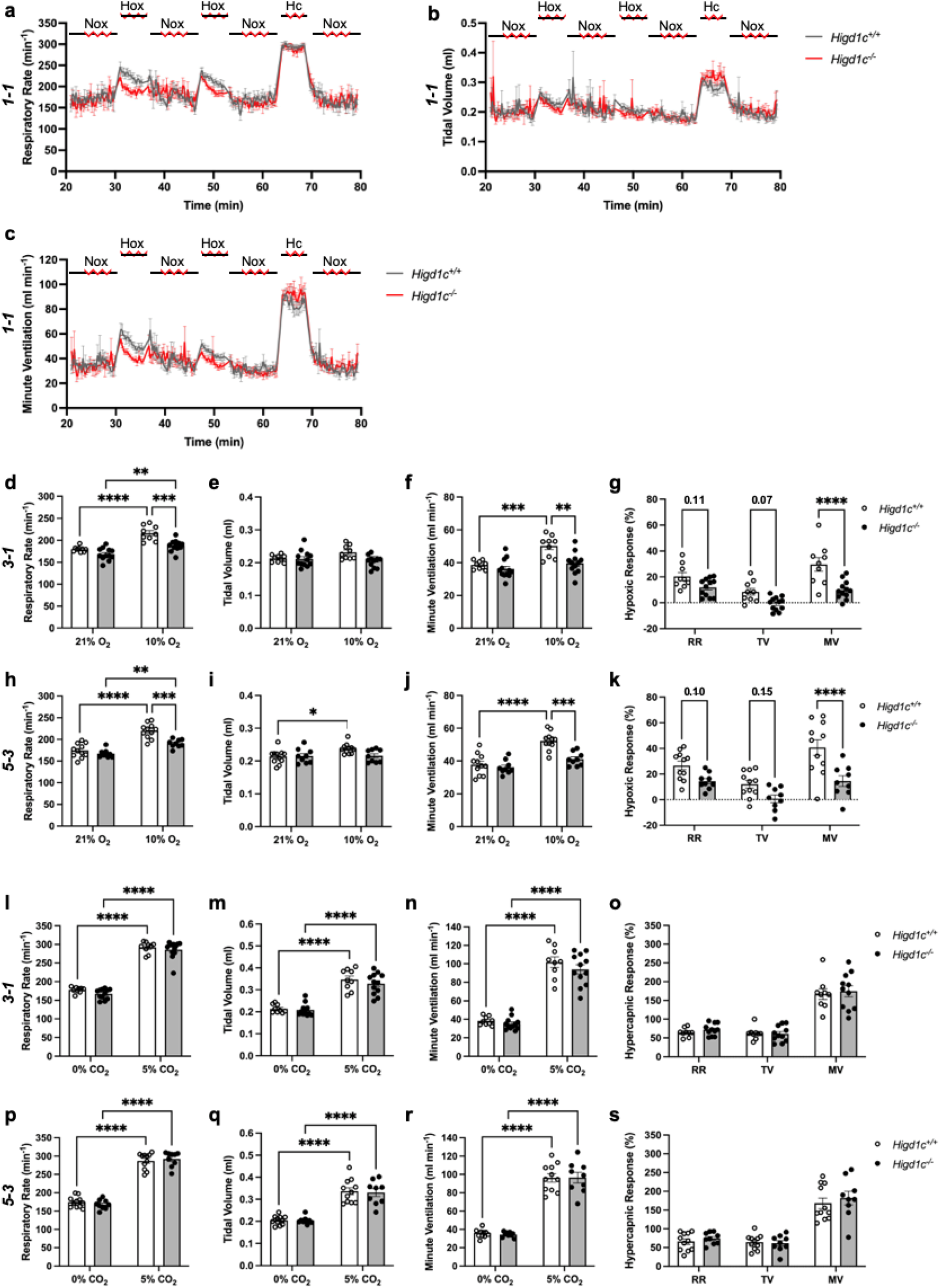
Ventilatory responses of *Higd1c 3-1* and *5-3* animals to hypoxia and hypercapnia. a-s, Respiratory rate (RR), tidal volume (TV), and minute ventilation (MV) (minute ventilation = respiratory rate × tidal volume) by whole-body plethysmography of unrestrained, unanesthetized *Higd1c^+/+^* and *Higd1c^-/-^* animals. a-c, Time course of experiments on *Higd1c 1-1* allele showing periods of normoxia (Nox), hypoxia (Hox), and hypercapnia (Hc). d-k, Ventilation of *Higd1c^+/+^* and *Higd1c^-/-^* animals of *3-1* (d-g) and *5-3* alleles (h-k) exposed to hypercapnia. g, k, Hypoxic response as percentage change in hypoxia (10% O_2_) versus control (21% O_2_). l-s, Ventilation of *Higd1c^+/+^* and *Higd1c^-/-^* animals of *3-1* (l-o) and *5-3* alleles (p-s) exposed to hypercapnia. o, s, Hypercapnic response as percentage change in hypercapnia (5% CO_2_) versus control (0% CO_2_). Ventilatory parameters of *Higd1c^+/+^* and *Higd1c^-/-^* animals in normal air conditions (21% O_2_ or 0% CO_2_) are not significantly different (p>0.05). n=11 (*1-1^+/+^*), 11 (*1-1^-/-^*), 9 (*3-1^+/+^*), 12 (*3-1^-/-^*), 11 (*5-3^+/+^*), 9 (*5-3^-/-^*) animals. Data as mean ± SEM. *p<0.05, **p<0.01, ***p<0.001, ****p<0.0001 by two-way ANOVA with Tukey’s test (d-f, h-j, l-n, p-r) or with Sidak correction (g, k, o, s).

**Figure S6.**
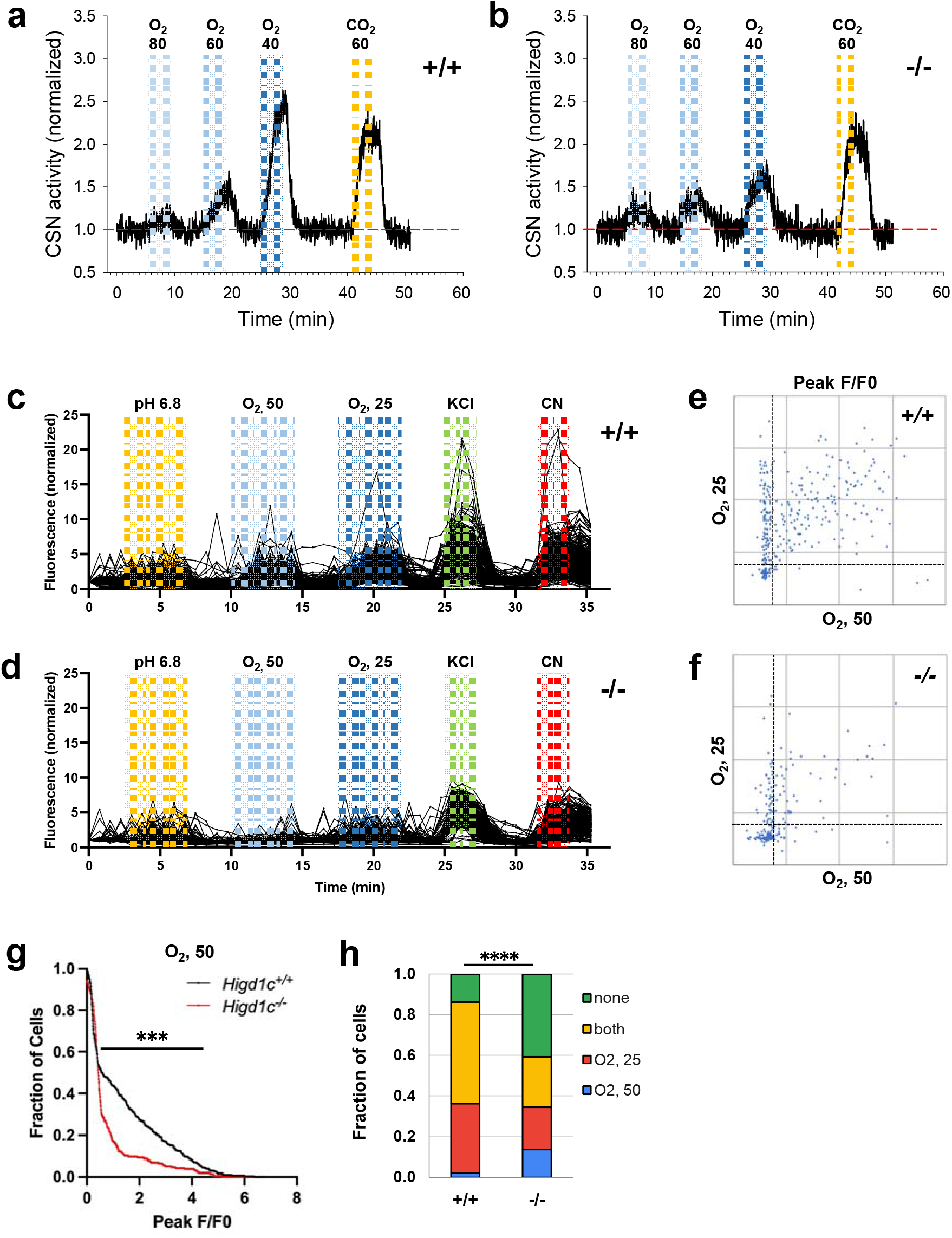
Calcium responses of *Higd1c 1-1* glomus cells to sensory stimuli. a, b, Representative traces of carotid sinus nerve (CSN) activity from *Higd1c 1-1^+/+^* and *Higd1c 1-1^-/-^* tissue preparations exposed to hypoxia (PO_2_=80, 60, and 40 mmHg) or hypercapnia (PCO_2_=60 mmHg) normalized to activity in impulses/s at t=0. c, d, Individual traces of GCaMP fluorescence of glomus cells from *Higd1c^+/+^* and *Higd1c^-/-^* animals, normalized to fluorescence at t=0 s. Z stacks collected every 45 s. e, f, Scatter plot of peak responses individual glomus cells to two levels of hypoxia (PO_2_=50 and 25 mmHg) in CBs from *Higd1c^+/+^* (e) and *Higd1c^-/-^* (f) animals. Strong responders have F/F0>1.5 as delineated by dashed lines. g, Fraction of cells responding to mild hypoxia (PO_2_=50 mmHg) with peak F/F0 above the threshold on the x-axis. ***p<0.001 by Z test of proportions. h, Fraction of cells that respond to hypoxia (PO_2_=50 and/or 25 mmHg) as subdivided in e and f. ****p<0.0001 by Chi-square test. n=296/4/3 (+/+), 201/4/3 (-/-). n as cells/CBs/animals.

**Figure S7.**
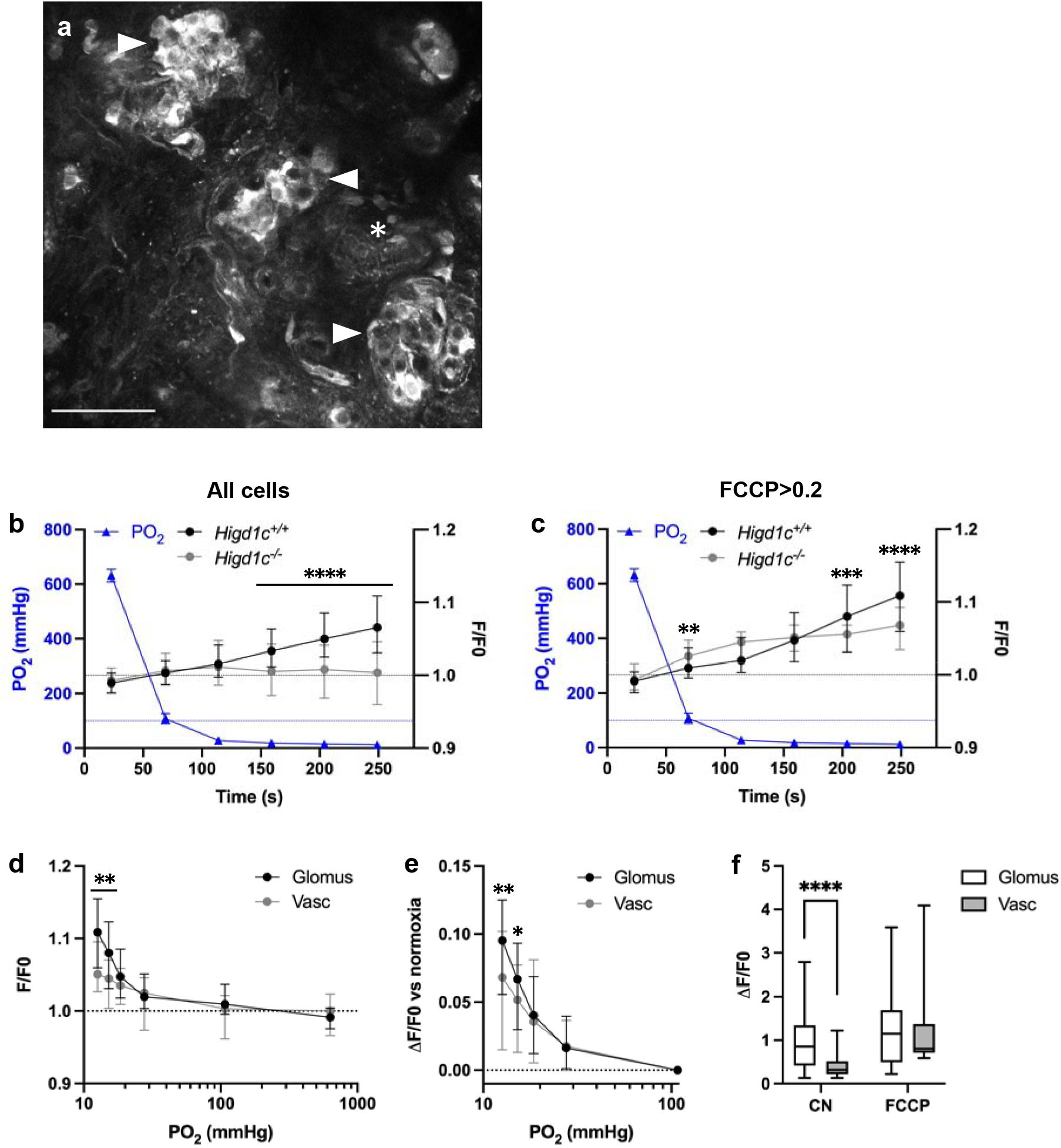
HIGD1C regulates the hypoxic response of the electron transport chain in carotid body glomus cells. a, Fluorescence of a whole-mount carotid body loaded with rhodamine 123 (Rh123), a dye sensitive to changes in mitochondrial inner membrane potential, under quenching conditions. Arrowheads, glomus cell clusters. Asterisk, vasculature. Scale bar, 50 µm. b-f, Change in Rh123 fluorescence of cells in *Higd1c 1-1^+/+^* and *Higd1c 1-1^-/-^* CBs in response to hypoxia (PO_2_<80 mmHg), cyanide (1 mM), and FCCP (8 µM). b, c, Time course of fluorescence relative to start of hypoxia stimulus. Data from all glomus cells (b) or only glomus cells with responses to FFCP at ΔF/F>0.2 (c-f). All vascular cells analyzed responded to FCCP with ΔF/F>0.2. b-c, Dashed blue line, normoxic PO_2_=100 mmHg comparable to well-oxygenated arterial blood. b-d, Dashed black line, fluorescence at the start of the stimulus. e, Dashed black line, ΔF/F at normoxia. b, n=291/3/3 (+/+), 312/3/3 (-/-) glomus cells/CBs/animals. c-f, n=98/3/3 (+/+), 102/3/3 (-/-) glomus cells/CBs/animals. d-f, 49/3/3 (+/+) vascular cells/CBs/animals. b-e, Data as median and interquartile interval. f, Data as box plots showing median, interquartile interval, and max/min. b-f, **p<0.01, ***p<0.001, ****p<0.0001 by Mann-Whitney U test with Holm-Sidak correction.

**Figure S8.**
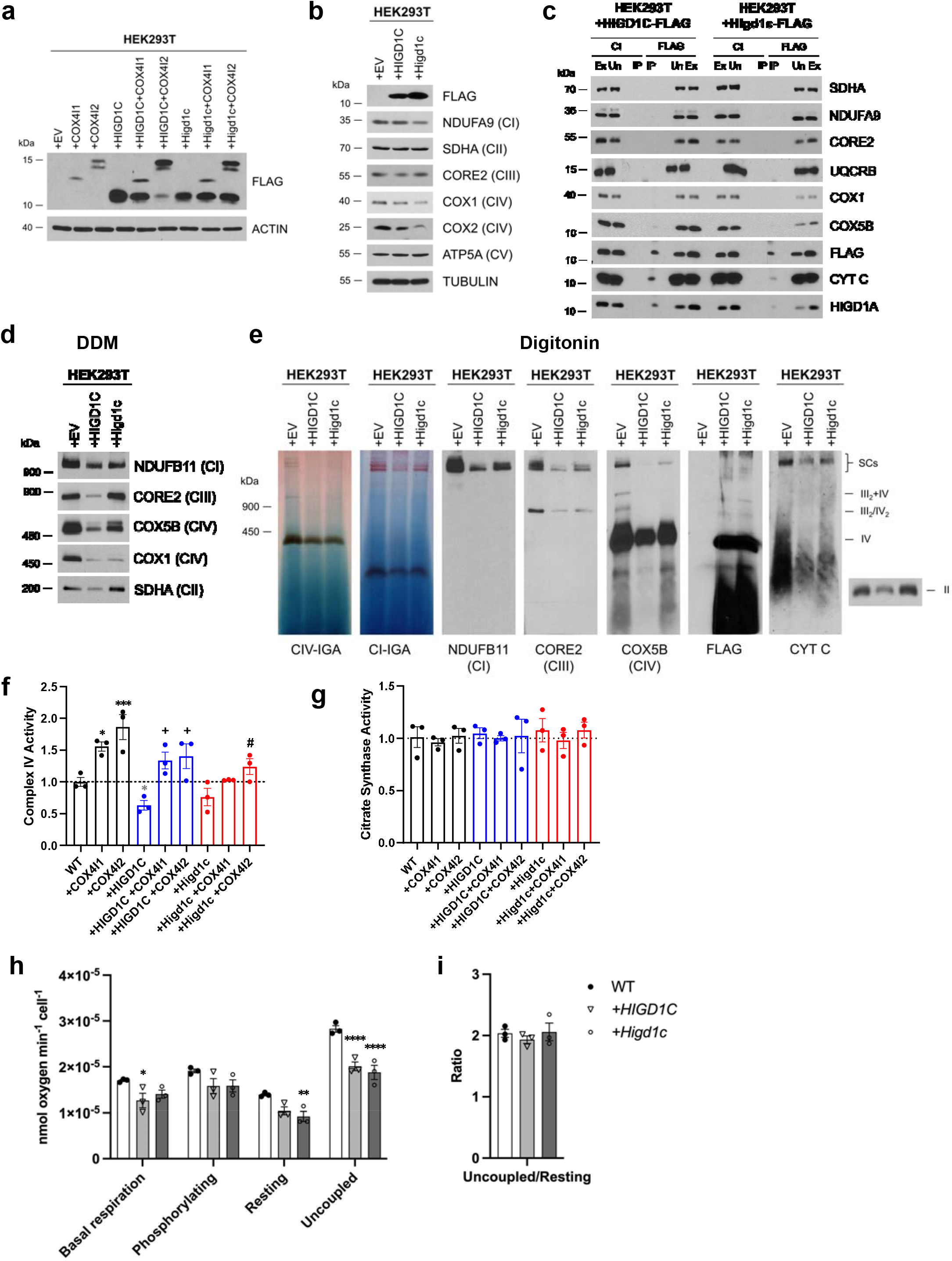
HIGD1C overexpression in wild-type HEK293T cells inhibits CIV activity. a-f, HEK293T cells overexpressing FLAG-tagged human or mouse HIGD1C or/and COX4I1/I2 isoforms. EV, empty vector. HIGD1C, human HIGD1C. Higd1c, mouse HIGD1C. a, SDS-PAGE and immunoblot using FLAG antibody to detect overexpression of different proteins. b, SDS-PAGE and immunoblots using antibodies for FLAG and ETC subunits. c, Co-immunoprecipitation using a FLAG antibody followed by SDS-PAGE and immunoblotting. d, BN-PAGE and immunoblotting from purified mitochondria extracted with DDM to analyze ETC complexes. e, Mitochondrial extracts solubilized with digitonin and analyzed by immunoblotting against antibodies for ETC complexes I, II, III and IV or by *in-gel* activity assays. f-g, Spectrophotometric measurement of CIV (f) or citrate synthase (g) activities in whole-cell extracts. Enzyme activities are expressed as a fraction of WT values. n=3. Data as mean ± SEM. *p<0.05, ***p<0.001 by one-way ANOVA with Dunnett test. *p-value versus WT, +p-value versus WT+HIGD1C, #p-value versus WT+Higd1c. h, Polarographic assessment in digitonin-permeabilized cells of KCN-sensitive oxygen consumption driven by succinate and glycerol-3-phosphate, in the presence or absence of ADP, oligomycin, and the uncoupler CCCP. n=3. i, Respiratory control ratio of the measurements performed in h. Data as mean ± SEM. *p<0.05, **p<0.01, ****p<0.0001 by one-way ANOVA with Dunnett test compared to WT.

**Figure S9.**
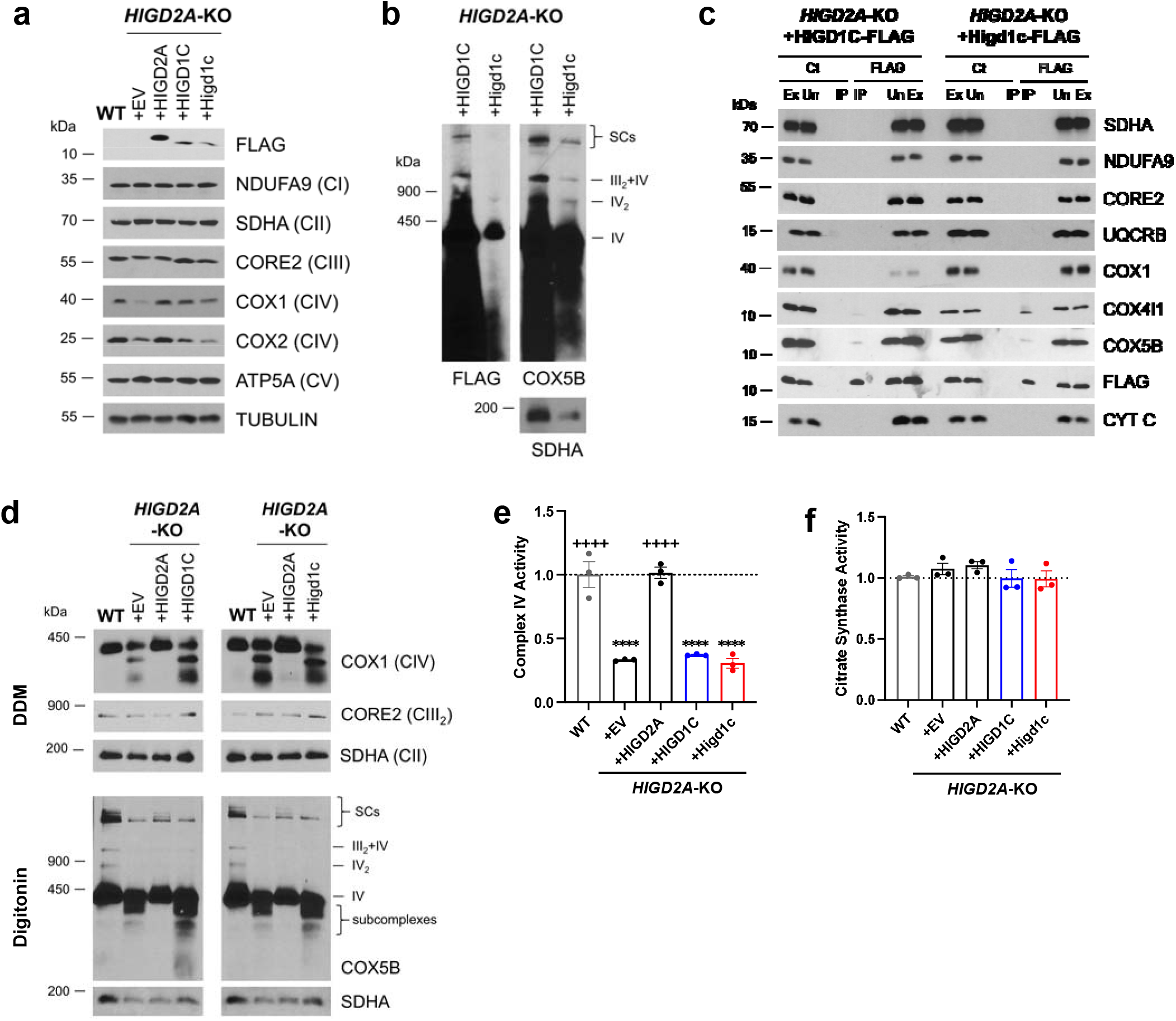
HIGD1C overexpression in *HIGD2A*-KO HEK293T cells does not rescue ETC defects. a-f, *HIGD2A-*KO cells overexpressing FLAG-tagged human or mouse HIGD1C. EV, empty vector. HIGD1C, human HIGD1C. Higd1c, mouse HIGD1C. a, SDS-PAGE and immunoblots using FLAG and ETC subunits. b, BN-PAGE of digitonin-permeabilized purified mitochondria and immunoblot using FLAG and COX5B antibodies, and SDHA as a loading control. c, Co-immunoprecipitation using a FLAG antibody followed by SDS-PAGE and immunoblotting. d, BN-PAGE and immunoblotting using purified mitochondria solubilized with DDM or digitonin, to analyze ETC complexes and supercomplexes, respectively. e-f, CIV and citrate synthase activities analyzed by polarography in whole-cell extracts. Enzyme activities are expressed as a fraction of WT values. n=3. Data as mean ± SEM. ****p<0.0001 by one-way ANOVA with Dunnett test. ****p<0.0001 versus WT, ++++p<0.0001 versus *HIGD2A*-KO+EV.

**Figure S10.**
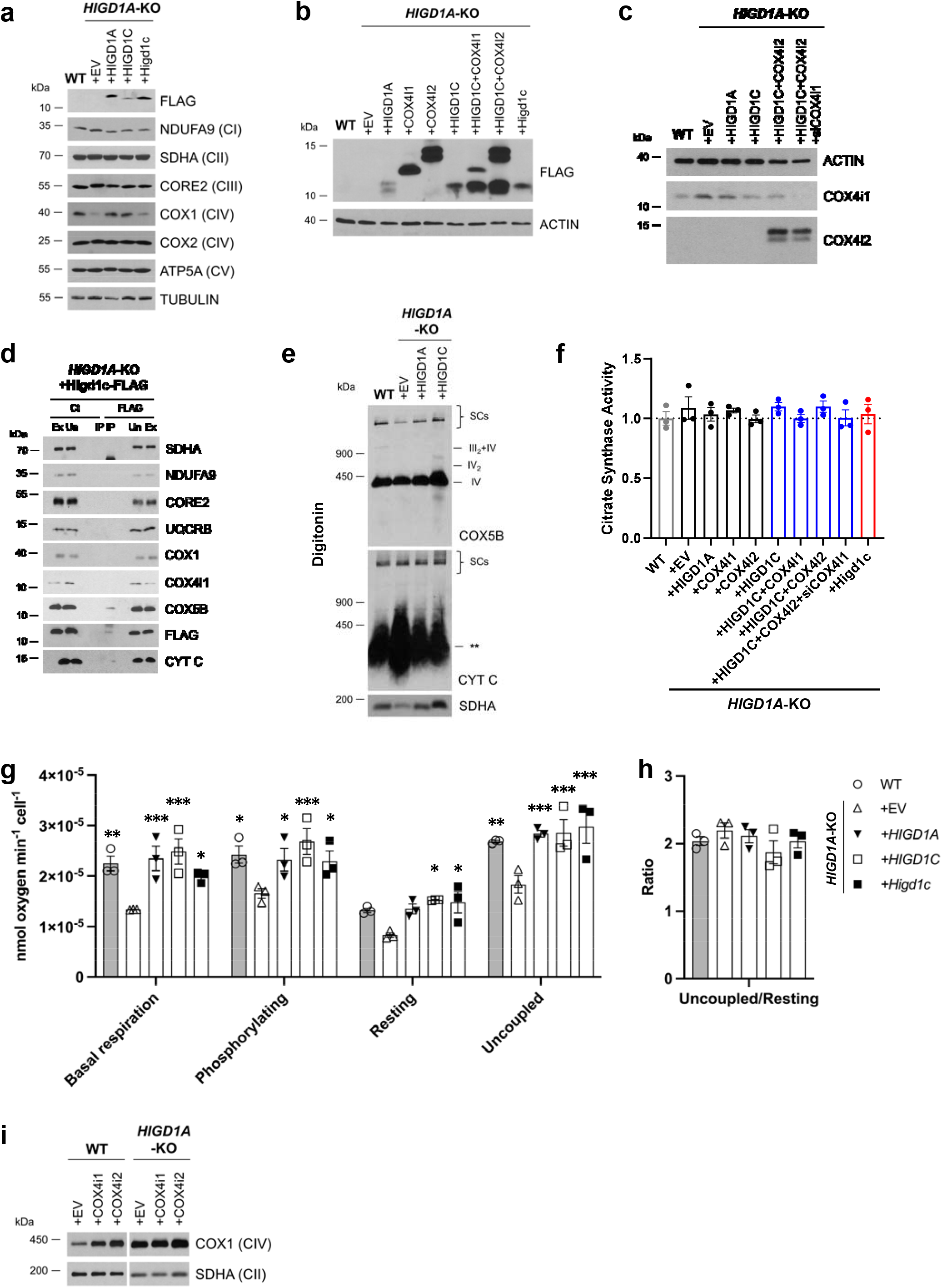
HIGD1C associates with CIV in *HIGD1A*-KO HEK293T cells and restores some ETC functions mediated by HIGD1A. a-f, *HIGD1A-*KO cells overexpressing FLAG-tagged human or mouse HIGD1C or/and COX4I1/I2 isoforms. EV, empty vector. HIGD1C, human HIGD1C. Higd1c, mouse HIGD1C. a, SDS-PAGE and immunoblots detecting FLAG and ETC subunits. b, SDS-PAGE and immunoblot using FLAG antibody to detect overexpression of the different proteins. c, SDS-PAGE and immunoblot using COX4I1 and COX4I2 antibodies to detect gene knockdown and overexpression, respectively. d, Co-immunoprecipitation using a FLAG antibody followed by SDS-PAGE and immunoblotting. e, BN-PAGE and immunoblotting using purified mitochondria solubilized with DDM or digitonin, to analyze ETC complexes and supercomplexes, respectively. ** indicates the accumulation of cytochrome *c* in an unidentified subcomplex. f, Citrate synthase activity measured by polarography in whole-cell extracts and expressed as a fraction of WT values. n=3. g, Polarographic assessment in digitonin-permeabilized cells of KCN-sensitive oxygen consumption driven by succinate and glycerol-3-phosphate, in the presence or absence of ADP (basal respiration and phosphorylating), oligomycin (resting), and the uncoupler CCCP (uncoupled). n=3. Data as mean ± SEM. *p<0.05, **p<0.01, ***p<0.001 vs. *HIGD1A-*KO +EV by two-way ANOVA with Dunnett test. h, Respiratory control ratio of the measurements performed in f. i, BN-PAGE and immunoblotting using digitonin-solubilized whole cells treated with DDM to analyze the levels of CIV. SDHA is used as a loading control.

**Figure S11.**
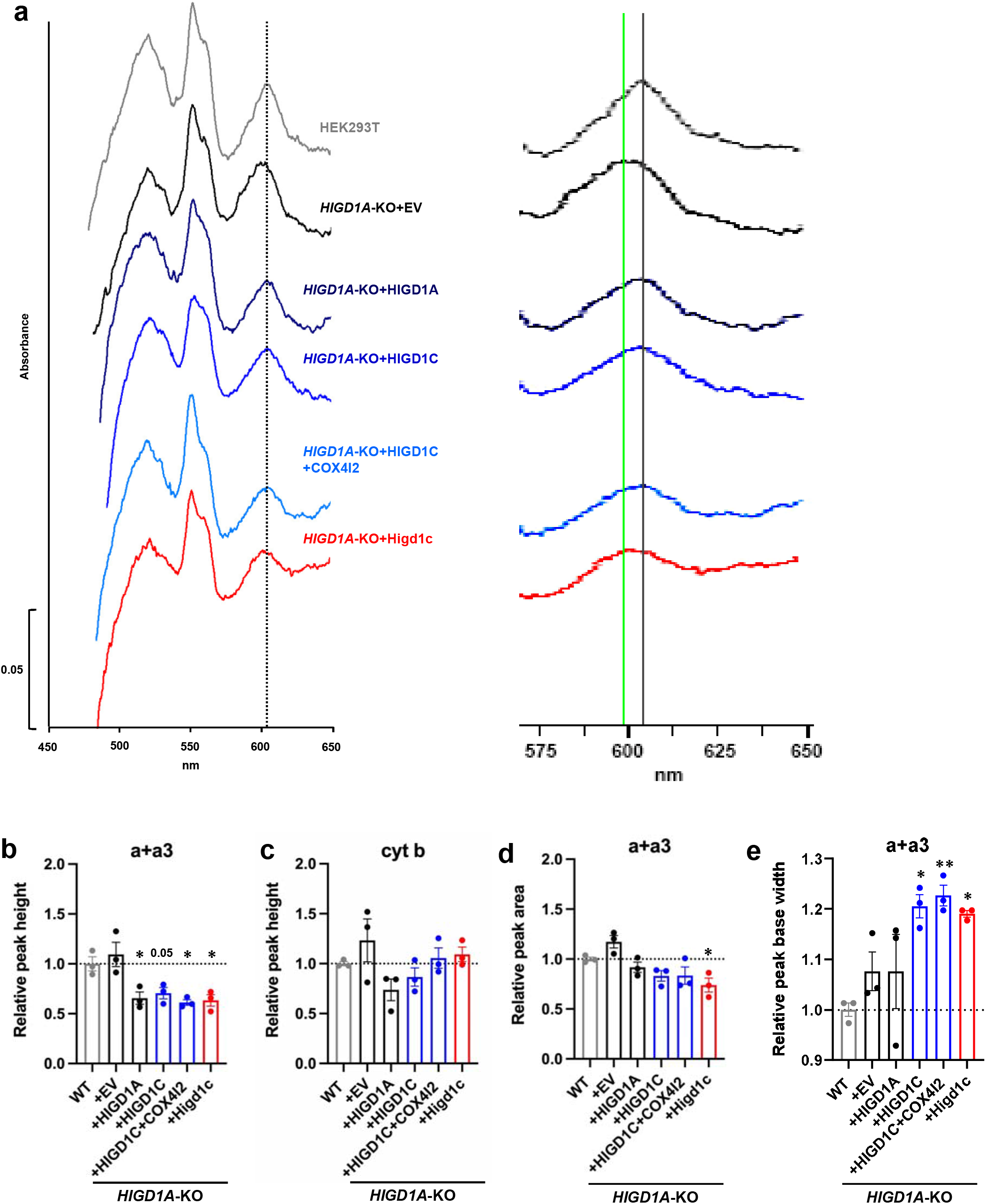
HIGD1C overexpression alters the oxygen binding domain of Complex IV in *HIGD1A*-KO HEK293T cells. a, Differential spectra (reduced minus oxidized) of total mitochondrial cytochromes measured by spectrophotometry. Absorbance of cytochromes extracted from purified mitochondria was measured from 450-650 nm. The panel on the left is a 2.5X magnification on the x-axis of the indicated portion of the graph. The grey vertical line indicates the wild-type heme a+a_3_ peak at 603 nm. The green vertical line marks the 599 nm wavelength, towards which some of the heme a+a_3_ peaks are shifted. b-c, Relative peak height of a+a3 (b) or cytochrome b (c) shown as ratio of WT. d, Relative peak area a+a3 measured as ratio of WT. e, Relative peak base width of a+a3 represented as ratio of WT. b-e, n=3. Data as mean ± SEM. *p<0.05, **p<0.01 by one-way ANOVA with Dunnett test.

**Table S1.**
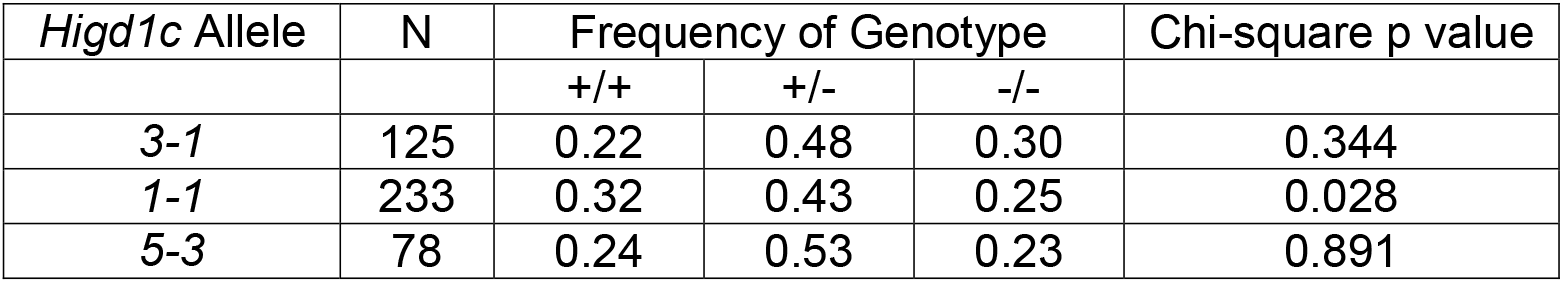
Genotype distribution of progeny from *Higd1c*^+/-^ x *Higd1c*^+/-^ crosses

| *Higd1c* Allele | N | Frequency of Genotype |  |  | Chi-square p value |
| --- | --- | --- | --- | --- | --- |
|  |  | +/+ | +/- | -/- |  |
| 3-1 | 125 | 0.22 | 0.48 | 0.30 | 0.344 |
| 1-1 | 233 | 0.32 | 0.43 | 0.25 | 0.028 |
| 5-3 | 78 | 0.24 | 0.53 | 0.23 | 0.891 |

**Table S2.**
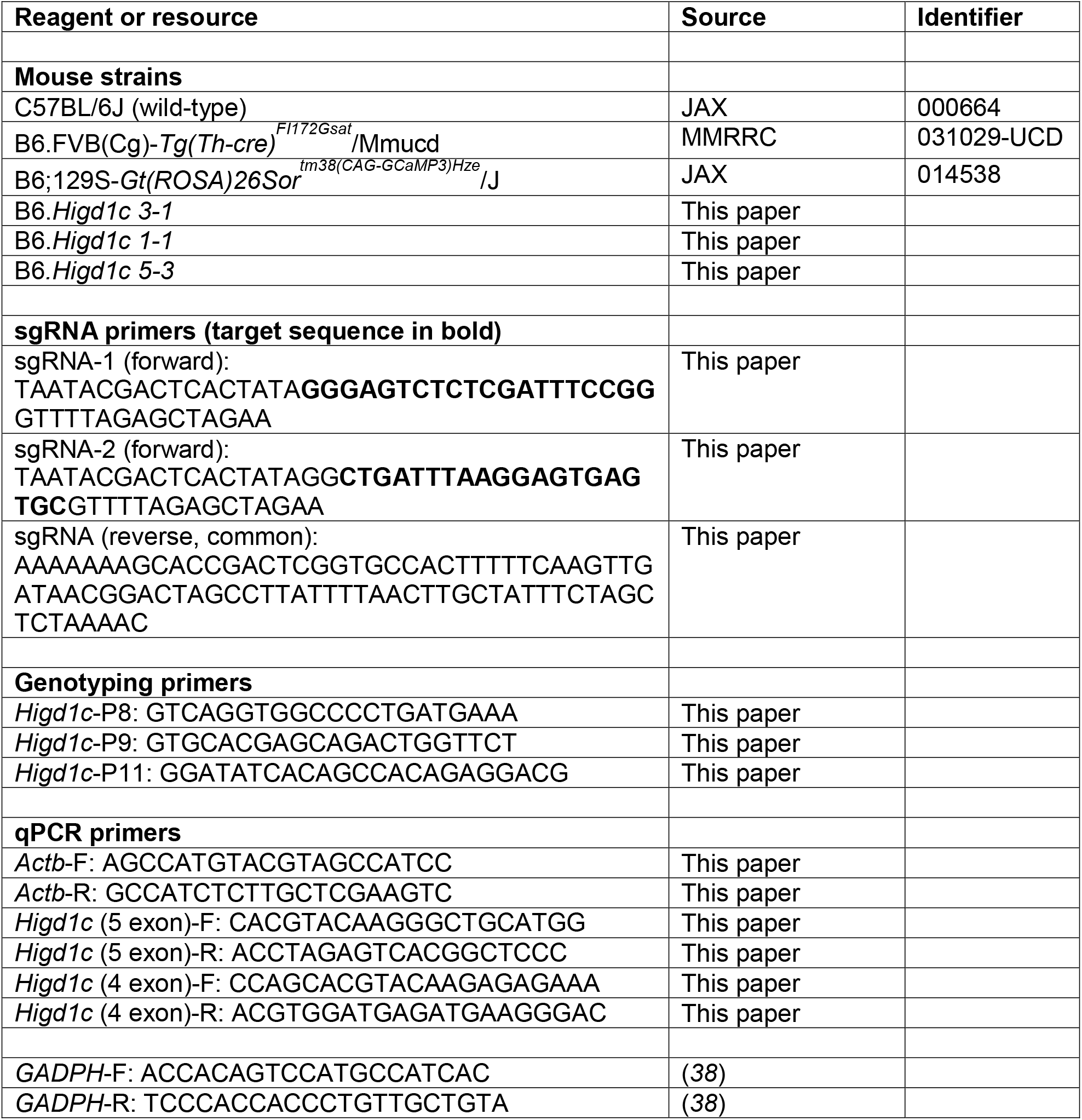
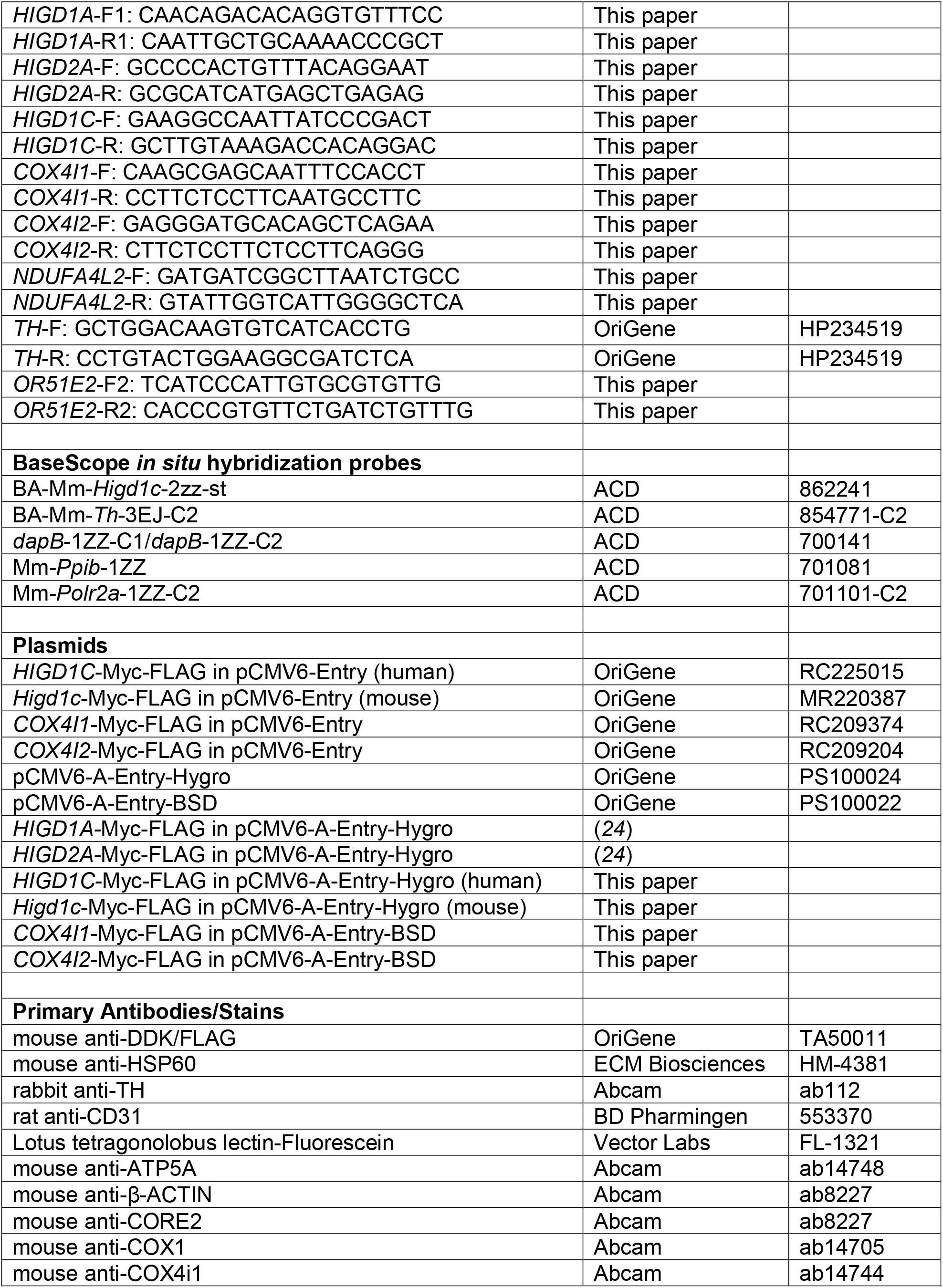
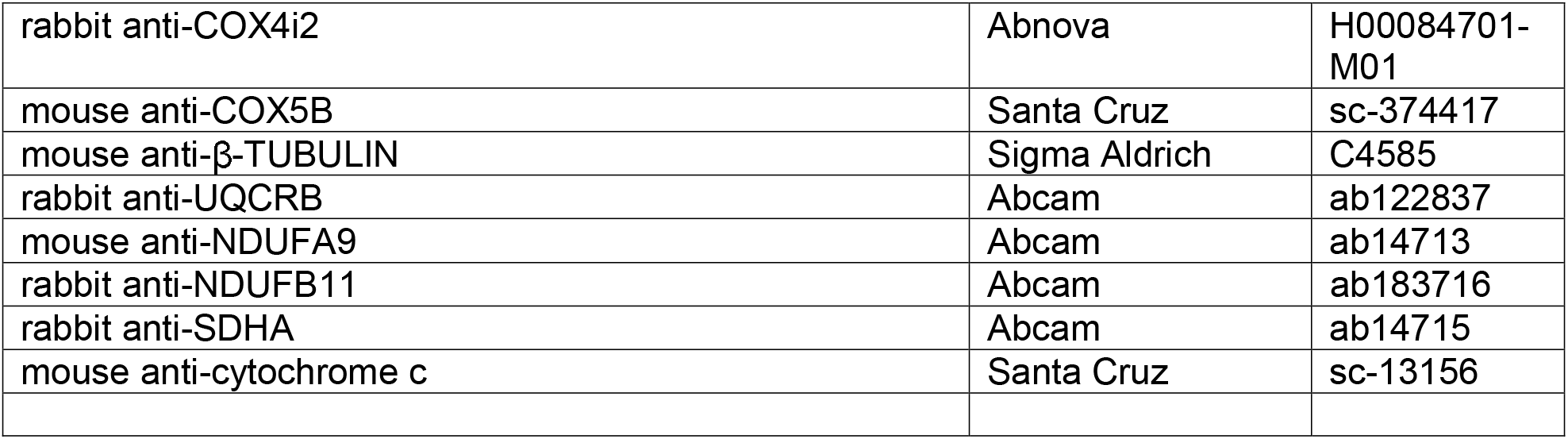
Reagents and resources

